# ACAD10 and ACAD11 enable mammalian 4-hydroxy acid lipid catabolism

**DOI:** 10.1101/2024.01.09.574893

**Authors:** Edrees H. Rashan, Abigail K. Bartlett, Daven B. Khana, Jingying Zhang, Raghav Jain, Andrew J. Smith, Zakery N. Baker, Taylor Cook, Alana Caldwell, Autumn R. Chevalier, Brian F. Pfleger, Peng Yuan, Daniel Amador-Noguez, Judith A. Simcox, David J. Pagliarini

**Affiliations:** Department of Biochemistry, University of Wisconsin–Madison, Madison, WI 53706, USA; Morgridge Institute for Research, Madison, WI 53715, USA; Department of Cell Biology and Physiology, Washington University School of Medicine, St. Louis, Missouri 63110, USA; Department of Biochemistry and Molecular Biophysics, Washington University School of Medicine, St. Louis, Missouri 63110, USA; Department of Genetics, Washington University School of Medicine, St. Louis, Missouri 63110, USA; Department of Microbiology, University of Wisconsin–Madison, Madison, WI 53706, USA; Department of Chemical and Biological Engineering, University of Wisconsin–Madison, Madison, WI 53706, USA; Department of Pharmacological Sciences, Icahn School of Medicine at Mount Sinai, New York, NY 10029 USA

**Author notes:** These authors contributed equally (Edrees H. Rashan, Abigail K. Bartlett).

## Abstract

Fatty acid β-oxidation (FAO) is a central catabolic pathway with broad implications for organismal health. However, various fatty acids are largely incompatible with standard FAO machinery until they are modified by other enzymes. Included among these are the 4-hydroxy acids (4-HAs)—fatty acids hydroxylated at the 4 (γ) position—which can be provided from dietary intake, lipid peroxidation, and certain drugs of abuse. Here, we reveal that two atypical and poorly characterized acyl-CoA dehydrogenases (ACADs), ACAD10 and ACAD11, drive 4-HA catabolism in mice. Unlike other ACADs, ACAD10 and ACAD11 feature kinase domains N-terminal to their ACAD domains that phosphorylate the 4-OH position as a requisite step in the conversion of 4-hydroxyacyl-CoAs into 2-enoyl-CoAs—conventional FAO intermediates. Our ACAD11 cryo-EM structure and molecular modeling reveal a unique binding pocket capable of accommodating this phosphorylated intermediate. We further show that ACAD10 is mitochondrial and necessary for catabolizing shorter-chain 4-HAs, whereas ACAD11 is peroxisomal and enables longer-chain 4-HA catabolism. Mice lacking ACAD11 accumulate 4-HAs in their plasma while comparable 3- and 5-hydroxy acids remain unchanged. Collectively, this work defines ACAD10 and ACAD11 as the primary gatekeepers of mammalian 4-HA catabolism and sets the stage for broader investigations into the ramifications of aberrant 4-HA metabolism in human health and disease.

## Main

The core enzymes of mitochondrial fatty acid β-oxidation (FAO) were discovered in the mid-20^th^ century following the classic cell fractionation work of Claude, Lehninger, and others^1-3^. In the ensuing decades, additional enzymes (e.g., reductases and isomerases) required to process select unsaturated fatty acids before they could enter the FAO cycle were identified. Additionally, a second venue for FAO—the peroxisome—was revealed, along with enzymes in this organelle critical for the catabolism of branched and very long-chain fatty acids^4,5^. To date, at least 17 proteins related to fatty acid catabolism have been directly linked to human genetic diseases, and aberrant fatty acid metabolism is associated with many metabolic disorders, including obesity, diabetes, glioblastoma, and chronic liver and kidney dysfunction^6-11^.

Discoveries of new fatty acid species—driven by advances in analytical techniques, such as mass spectrometry (MS)^12^—are further expanding our understanding of lipid metabolism and highlighting the requirement for additional enzymes that enable their production and catabolism. A key recent example is the family of branched **f**atty **a**cyl esters of **h**ydroxy **f**atty **a**cids (FAHFAs), which has been linked to beneficial effects on glycemia, insulin secretion, and inflammation^13^. A second, less studied family is the 4-hydroxy acids (4-HAs), which comprises fatty acids with a hydroxy (–OH) group on the 4 (γ) position. 4-HAs are produced by common lipid peroxidation, formed by enzymatic conversion or the catabolism of longer-chain hydroxy acids, and can be ingested in the form of various naturally-occurring lipids or certain drugs of abuse (e.g. 4-hydroxybutyrate)^14-17^. Indeed, recent mass spectrometry analyses have identified a spectrum of 4-HAs and 4-HA precursors in human serum, yet few 4-HAs have been studied at any detail in mammalian systems^12^.

### ACAD10/11 process 4-HA substrates *in vitro* and in cells

A central question in 4-HA biology is what enzymes catabolize these lipids following their acylation with coenzyme A (CoA). The presence of an –OH group on the 4^th^ (γ) carbon likely reduces their compatibility with the established acyl-CoA dehydrogenases (ACADs) of FAO, which catalyze the α,β-dehydrogenation of acyl-CoA substrates to produce 2-enoyl-CoAs (Fig. 1a). Nonetheless, previous MS-based metabolic tracing analyses performed on murine livers perfused with various 4-HAs demonstrated that these lipids can be catabolized via two pathways: a minor pathway, involving a sequence of β-oxidation, α-oxidation, and β-oxidation steps; and a major pathway—5-6-fold more active than the minor—involving a unique phosphorylated acyl-CoA intermediate (Fig. 1a). However, no mammalian enzymes required for this major pathway have been identified^18-21^. More recently, a similar pathway was observed for the breakdown of levulinic acid (Lva) in *Pseudomonas putida*^22^. Here, Lva is converted into 4-hydroxyvalerate (4-HV), a short-chain 4-HA, that then proceeds into FAO via a similar phosphorylated intermediate. The authors further identified the necessary “*lva* operon” for this pathway, which includes genes encoding a kinase-like protein (LvaA) and an ACAD family protein (LvaC) (Fig. 1b).

**Fig 1.**
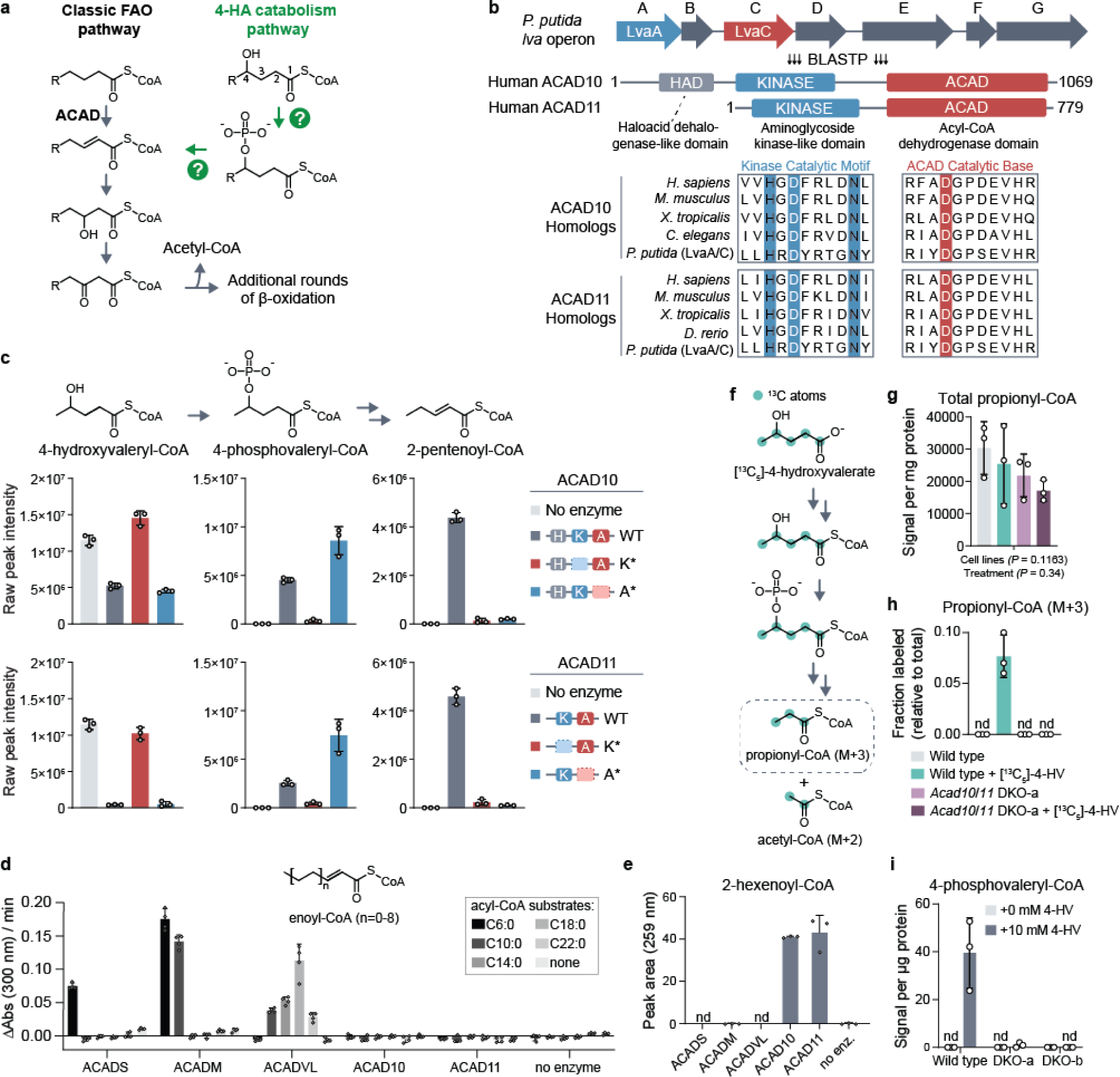
ACAD10/11 process 4-HA substrates *in vitro* and in cells. **(a)** A model of the major 4-hydroxy acid (4-HA) catabolism pathway and its integration into the classic fatty acid β-oxidation pathway based on findings of Zhang et al., 2009 and Harris et al., 2011. Question marks indicate uncharacterized enzymatic steps in mammals. **(b)** Multiple sequence alignment demonstrating conservation between LvaA and LvaC and eukaryotic ACAD10/11 homologs. Blue highlighted residues represent invariant residues of the kinase catalytic motif HxDx_4_N. Red highlighted residue represents the conserved, putative catalytic aspartate of the ACAD domain. **(c)** LC-MS analysis of *in vitro* enzyme reactions. 4-hydroxyvaleryl-CoA substrate was created by LvaE. Displayed are the raw signal intensities of 4-hydroxyvaleryl-CoA, 4-phosphovaleryl-CoA, and 2-pentenoyl-CoA with the corresponding chemical structures included above. Top and bottom sets of bar graphs represent data from reactions with recombinant ACAD10^NΔ34^ and ACAD11, respectively. The same “no enzyme” control data is displayed on both sets of bar graphs for clearer comparison. Data are expressed as mean -/+ SD (n = 3 replicate reactions). **(d)** Ferrocenium ion assay for ACADS, ACADM, ACADVL, ACAD10 and ACAD11 with unsubstituted acyl-CoA substrates. Rates correspond to the initial velocity of the enzyme reaction as measured by change in absorbance over time. The substrate lengths tested range from C6:0 to C22:0. The structure of the final product, enoyl-CoA, is shown (mean -/+ SD; n = 4 technical replicate reactions). **(e)** Quantification of 2-hexenoyl-CoA produced from 4-phosphohexanoyl-CoA by HPLC UV-Vis spectroscopy (mean -/+ SD; n = 3 technical replicate reactions). **(f)** Stable isotope tracing strategy to study 4-HV catabolism through the major pathway. **(g)** Total abundance of all propionyl-CoA isotopologs. Data represents the averaged sum of raw signal intensities of each isotopolog. Two-way ANOVA was used to test statistical significance. **(h)** Fractional abundance of M+3 propionyl-CoA relative to total propionyl-CoA in wild type or *Acad10*/*Acad11* DKO-a cells cultured with 5 mM [^13^C_5_]-4-HV. **(i)** 4-phosphovaleryl-CoA abundance detected in wild-type and DKO Hepa1-6 cells supplemented with 10 mM unlabeled 4-HV. Metabolomic results are expressed as mean -/+ SD; n = 3 replicate experiments.

To identify potential mammalian orthologs of LvaA and LvaC, we performed a BLASTP analysis. Surprisingly, the top hits for LvaA and LvaC were the same two proteins—ACAD10 and ACAD11 (ACAD10/11 when referring to both), two poorly characterized acyl-CoA dehydrogenase paralogs. LvaA exhibits high homology to N-terminal kinase domains found in ACAD10/11 but not in other members of the ACAD family (Fig. 1b; Extended Data Fig. 1a). These kinases possess all the core catalytic residues and subdomain motifs of the protein kinase-like (PKL) superfamily and show particular homology to the aminoglycoside kinase/phosphotransferase 3 (APH3) subgroup of eukaryotic-like kinases (Extended Data Fig. 4b)^23^. LvaC exhibited high homology to the C-terminal ACAD domains of ACAD10/11. Among all mammalian ACADs, LvaC is most similar to ACAD10/11 (Extended Data Fig. 1b) as it shares a predicted catalytic aspartate and other active site sequences only conserved between ACAD10/11 homologs. ACAD10 also has a haloacid dehalogenase-like domain (Fig. 1b) that lacks homology to any protein in the *lva* operon and was not evaluated further here. Overall, our sequence conservation analysis supports ACAD10/11 as bifunctional homologs of LvaA and LvaC and candidates for the missing enzymes of mammalian 4-HA catabolism.

We next purified recombinant wild-type and two mutant versions of ACAD10 and ACAD11 to test their *in vitro* activities against a 4-hydroxyacyl-CoA substrate (Extended Data Fig. 1c,d). In the first mutant (K*), we replaced the catalytic aspartate of the kinase domain with alanine, thereby eliminating kinase activity but leaving ACAD activity intact. In the second mutant (A*), we replaced the catalytic aspartate of the ACAD domain with asparagine, eliminating the ACAD activity but leaving kinase activity intact. We used recombinant LvaE, an acyl-CoA synthetase encoded in the *lva* operon^22^, to produce 4-hydroxyvaleryl-CoA (4-HV-CoA) as a substrate for *in vitro* reactions. The activities of ACAD10/11 were evaluated by measuring intermediates expected to originate from 4-HV-CoA using LC-MS and HPLC (Extended Data Fig. 1e). The wild-type version of each enzyme converted 4-HV-CoA into 2-pentenoyl-CoA as a final product while also producing differing levels of the 4-phosphovaleryl-CoA (4-PV-CoA) intermediate (Fig. 1c; Extended Data Fig. 1g). The K* mutants demonstrated little to no substrate consumption, whereas the A* mutants generated the 4-PV-CoA intermediate but were unable to complete conversion into 2-pentenoyl-CoA. We also observe formation of 3-hydroxyvaleryl-CoA in our reactions but determine it to be a non-enzymatic byproduct of 2-pentenoyl-CoA (Extended Data Fig. 1f).

Our results demonstrate that ACAD10/11 are sufficient to convert 4-HV-CoA into 2-pentenoyl-CoA, a conventional FAO intermediate, thereby establishing them as the missing enzymes necessary to execute the major 4-HA catabolic route described previously^18^. Our results further demonstrate that the ACAD10/11 kinase reaction precedes the ACAD reaction, indicating that the ACAD domains of these enzymes must accommodate a phosphorylated substrate. Together, this implies that the ACAD10/11 ACAD domains are functionally distinct from other ACAD enzymes, consistent with evolutionary analyses demonstrating that ACAD10/11 share a more recent common ancestor with each other than with other ACADs^24^.

To test this hypothesis, we compared the activities of ACAD10/11 with three recombinantly purified ACAD enzymes involved in FAO: ACADS, ACADM, and ACADVL, which have well-established specificity for short-, medium-, and long/very long-chain acyl-CoA substrates, respectively. First, we tested the ability of these enzymes and ACAD10/11 to oxidize fatty acyl-CoAs ranging in length from 6-22 carbons into their 2-enoyl-CoA products using a ferrocenium assay^25^. ACADS, ACADM, and ACADVL each exhibited clear activity consistent with their established specificity (Fig. 1d). However, ACAD10/11 showed no activity against any substrate under these conditions. Previous work suggested that ACAD11 can accommodate unsubstituted longer-chain substrates similar to those tested here^26^. However, these experiments were performed using bacterial lysates expressing a truncated form of ACAD11. While it is possible that ACAD10/11 may retain partial activity against classical FAO substrates under different conditions, the head-to-head comparison here demonstrates that this is unlikely.

We next compared the activities of each enzyme against 4-phosphohexanoyl-CoA (4-PH-CoA), an intermediate that can be generated by the kinase domains of ACAD10/11 from 4-hydroxyhexanoyl-CoA. We produced this intermediate using the ACAD11 A* mutant, which cannot act upon the phospho-intermediate. Wild-type ACAD10/11 given this intermediate successfully converted it into the 2-hexenoyl-CoA product (Fig. 1e), indicating that this substrate does not need to be channeled to the ACAD domain from the kinase domain. However, none of the other ACADs showed any detectable activity against 4-PH-CoA. Together, these results demonstrate that the ACAD domains of ACAD10/11 are functionally distinct from other ACADs, and they likely do not contribute appreciably to the catabolism of conventional fatty acids.

Finally, we evaluated whether ACAD10/11 are necessary for the major 4-HV catabolic pathway by performing stable isotope tracing in Hepa1-6 cells. To do so, we cultured wild-type and CRISPR/Cas9-mediated *Acad10*/*Acad11* double knockout (DKO) cells with [^13^C_5_]-4-HV and measured both total and labeled (M+3) propionyl-CoA (Fig. 1f). M+3 labeling of propionyl-CoA is only expected if 4-HV is catabolized via the 4-PV-CoA intermediate^19^ (Extended Data Fig. 1h). Each cell line possessed comparable levels of total propionyl-CoA (Fig. 1g); however, only wild-type Hepa1-6 cells produced M+3 propionyl-CoA (Fig. 1h). Consistently, none of the distinctive 4-PV-CoA intermediate was observed in the DKO cells even when cultured with high amounts of 4-HV (Fig. 1i). Overall, these results demonstrate that ACAD10/11 are sufficient for 4-HA conversion into physiological FAO substrates *in vitro* and are necessary for the major 4-HA catabolic pathway in Hepa1-6 cells.

### Cryo-EM of ACAD11 reveals structural determinants of molecular function

To further explore the bifunctionality of ACAD10/11 at the molecular level, we initiated cryo-electron microscopy (cryo-EM) experiments with the ACAD11 K* and A* mutant proteins purified from *E. coli*. To explore the effect of substrate interaction on protein structure, these mutants were incubated with 4-HV-CoA and 4-PV-CoA, respectively, before preparation. Negative stain TEM analysis confirmed homogeneity and revealed monodispersed particles for each sample. The samples were vitrified, imaged, and reconstructed using a 3D single-particle cryo-EM analysis (Extended Data Fig. 2; Extended Data Fig. 3; Extended Data Table 1). A small number of the particles of the ACAD11 K* mutant revealed an intact structure, but only the ACAD domains were resolvable (Extended Data Fig. 4a). The tetrameric structure was consistent with other known ACADs, including an unpublished structure of a truncated human ACAD11 that included only the ACAD domain^27^ (PDB ID: 2WBI). However, both domains were resolved in the ACAD11 A* mutant structure, revealing ordered interactions between the kinase and ACAD domains (Fig. 2a).

**Fig 2.**
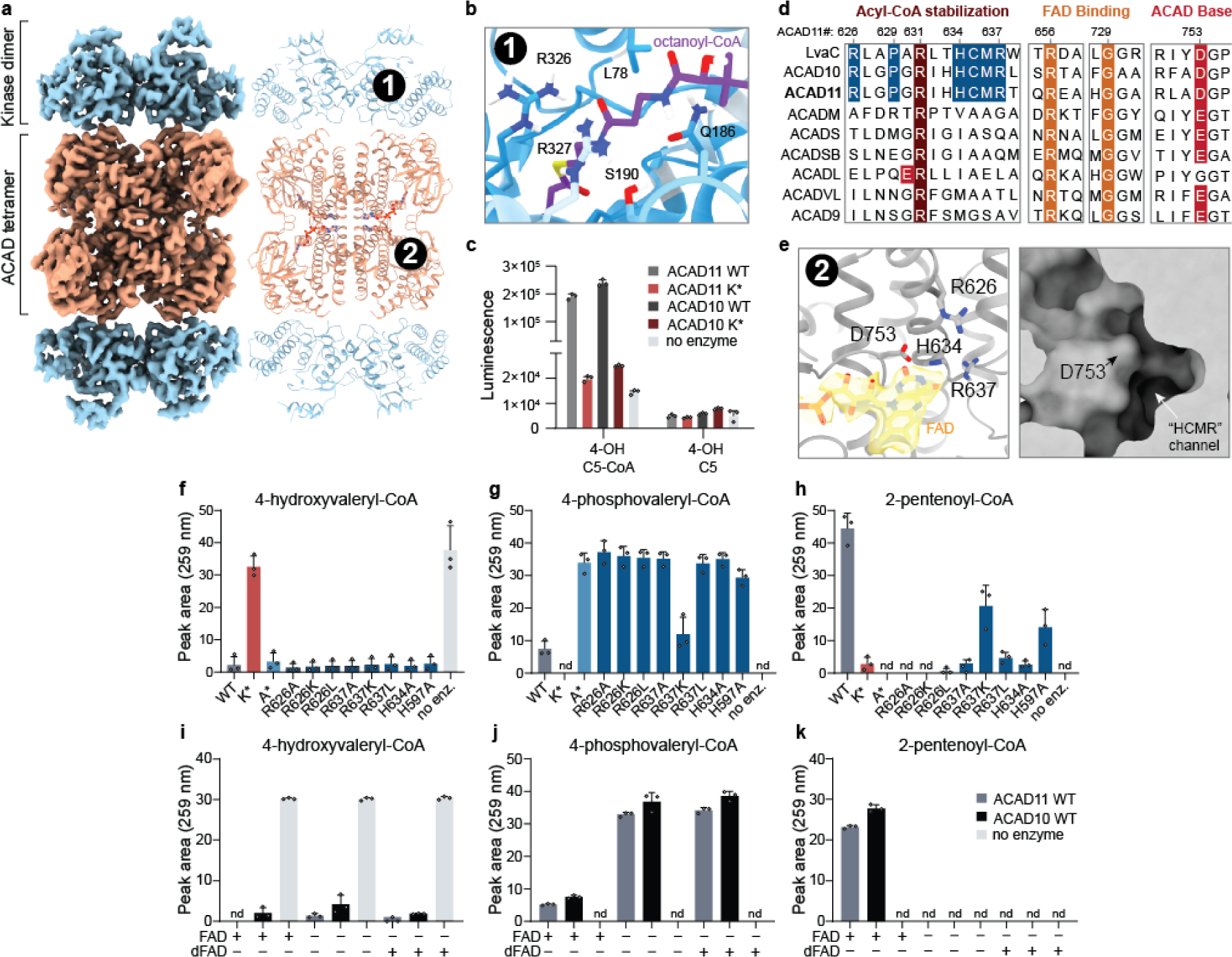
Cryo-EM of ACAD11 reveals structural determinants of molecular function. **(a)** Cryo-EM map and model of the ACAD11 A* mutant tetramer incubated with 4-PV-CoA. The kinase and ACAD domains are colored blue and orange respectively. **(b)** *Ab initio* HADDOCK docking of octanoyl-CoA into the kinase domain active site. The pantothenic acid region of the octanoyl-CoA ligand is centered (purple). CONSURF residue conservation scores are shown as a gradient from blue to white, with blue as most conserved for ACAD10/11 and LvaA homologs. Proposed interacting residues, L78, Q186, S190, R326, and R327, are presented as sticks. **(c)** ADP quantification for reaction mixtures containing ACAD10/11 wild type or mutant with 4-hydroxyvaleryl-CoA or 4-hydroxyvalerate substrate. **(d)** Sequence alignments for *M. musculus* ACAD10/11 with *P. putida* LvaC and other *M. musculus* ACAD family members. Key, highly conserved residues involved in acyl-CoA stabilization, FAD binding and catalytic activity are highlighted in red and orange. Residues proposed as potential ACAD10/11 functional markers are highlighted in blue. **(e)** FAD binding site in the structure of the ACAD11 K* mutant with 4-HV-CoA. The FAD model and map are highlighted (yellow). Critical residues in the active site, R626, R637, D753 and H634, are presented as sticks. A surface representation of the ACAD domain active site pocket is shown in the right panel. **(f-h)** UV-Vis HPLC quantification (259 nm) of 4-hydroxyvaleryl-CoA, 4-phosphovaleryl-CoA, and 2-pentenoyl-CoA in reactions containing ACAD11 wild type and mutants. **(i-k)** UV-Vis HPLC quantification (259 nm) of 4-hydroxyvaleryl-CoA, 4-phosphovaleryl-CoA, and 2-pentenoyl-CoA in reactions containing 5 µM FAD or 5-deazaFAD (dFAD). All assay data are represented as mean -/+ SD (n = 3 technical replicates).

The core ACAD11 kinase fold is similar to that of well-characterized PKLs^23,28,29^ (Extended Data Fig. 4b), whereby the N lobe consists of an extended β sheet and a single α helix (aC), and the C lobe is comprised of a series of α helices and β strands. The kinases domains form “back-to-back” dimers, with the active sites facing outward. Similar to other members of the APH3 family, the ACAD11 kinase includes a long insert between the αE helix and the catalytic subdomain that is typically important for substrate recognition. To understand how the kinase domain might bind an acyl-CoA substrate, we performed *ab initio* small molecule docking of octanoyl-CoA using HADDOCK^30,31^. While the carbon chain of octanoyl-CoA was positioned inside the active site pocket near the catalytic base (D220), the CoA moiety associated with the outer surface of the kinase domain. A series of residues that are highly conserved between LvaA and ACAD11 homologs line the opening of the kinase active site, suggesting interaction sites for the CoA group (Fig. 2b). Some of these hydrophilic residues, such as Q186 and R326, neighbor the pantothenic acid region of CoA; their interactions could support specificity of the enzyme for CoA-ligated substrates. Consistently, neither ACAD10 nor ACAD11 were able to phosphorylate the free acid form of 4-HV, indicating that the CoA moiety is necessary for substrate recognition (Fig. 2c). The kinase dimers also interact with the ACAD tetramer at interfaces mediated by a putative salt-bridge linkage, suggesting that a physical connection between the kinase and ACAD domains may assist efficient conversion of substrate to product. Together, these data demonstrate that the ACAD10/11 kinase domain is poised to recognize 4-hydroxylacyl-CoA species and generate a phosphorylated substrate for further processing by the ACAD domain.

The ACAD fold of our structure suggests that a set of amino acids conserved in ACAD10 and LvaC may explain ACAD11’s ability to accommodate a phosphorylated substrate (Fig. 2d). Within the structure, R626, H634, and R637 are found on a single α-helix within the active site that is directly adjacent to the catalytic aspartate D753. These residues point toward D753 and the open cavity of the active site. In addition, an atypical pocket situated directly between these residues is established by the highly conserved proline 629 (Fig. 2e). To investigate this region experimentally, we purified a series of ACAD10/11 constructs with mutations to these conserved residues, as well as to another histidine residue near the active site, and measured their activities against 4-HV-CoA (Extended Data Fig. 4c). As predicted, disruptions to the atypical pocket area ablated the activity of the ACAD domain, resulting in accumulation of the phospho-intermediate (Fig. 2f-h).

Our structure also exhibits density in the ACAD11 active site consistent with bound flavin adenine dinucleotide (FAD) (Fig. 2d,e). In typical ACADs, FAD is a requisite cofactor that is reduced to FADH_2_ concomitant with substrate oxidation^32^. However, the conversion of 4-PV-CoA to 2-pentenoyl-CoA does not involve a permanent change in oxidation state. To test whether ACAD10/11 require FAD for activity, we performed enzymatic assays on 4-HV-CoA with increasing concentrations of FAD, including sub-molar quantities, and quantified the substrate, intermediate, and product via HPLC. We observed that ACAD10/11 both require FAD to convert the phospho-intermediate into the final 2-enoyl-CoA product and that activity diminished as the FAD concentrations decreased (Extended Data Fig. 4d). To understand whether FAD is redox-active during this reaction, we substituted FAD with 5-deazaFAD, which limits the redox capacity of the cofactor while keeping binding and structural roles intact. ACAD10/11 showed no detectable ACAD domain activity in the presence of 5-deazaFAD, suggesting that these enzymes still require redox-active FAD to convert 4-phosphoacyl-CoA to 2-enoyl-CoA (Fig. 2i-k). This suggests that ACAD10/11 use FAD in an atypical manner distinct from other ACADs. Overall, these structure/function analyses reveal multiple distinctive features of ACAD10/11 that enable their processing of 4-HA-CoA substrates.

### ACAD10/11 mediate catabolism of distinct 4-HAs in different organelles

Our enzyme assays indicate that ACAD10/11 have comparable activities against the substrates tested *in vitro*; however, it is unclear whether they act redundantly in cellular 4-HA catabolism. Other acyl-CoA processing enzymes with similar inherent substrate specificities are known to reside in distinct subcellular locations where they encounter different substrates^33^. ACAD10 has a predicted N-terminal mitochondrial targeting sequence^34^ (MTS), and ACAD11 has a predicted C-terminal peroxisomal targeting sequence^35^ (PTS) (Fig. 3a); however, previous localization studies have provided conflicting results^26,36,37^.

**Fig 3.**
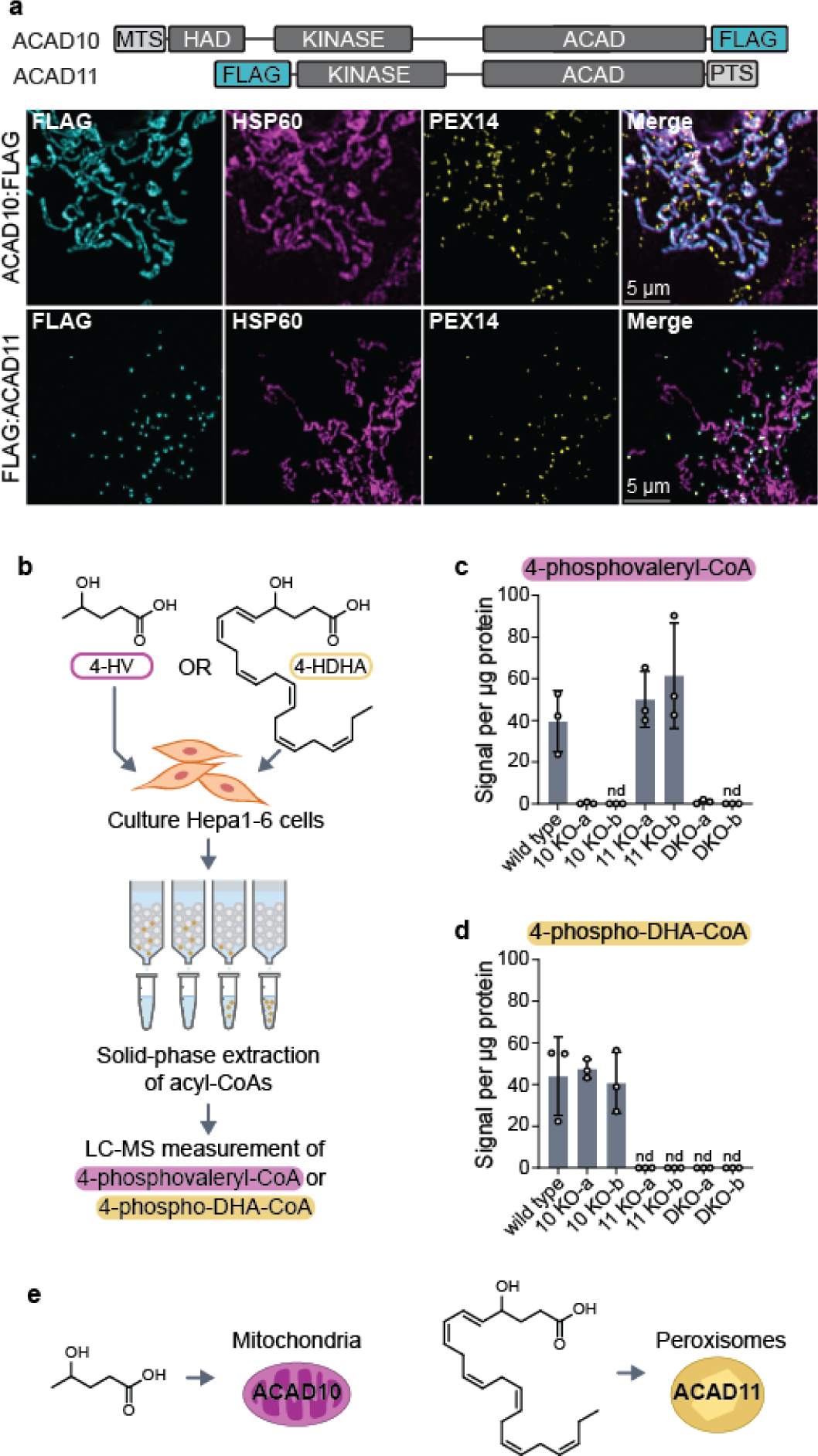
ACAD10/11 mediate catabolism of distinct 4-HAs in different organelles. **(a)** Immunofluorescence imaging of overexpressed human ACAD10:FLAG and FLAG:ACAD11 in COS7 cells. Localization of FLAG-tagged constructs (cyan) is indicated by overlap in fluorescence intensity with peroxisomal (PEX14; yellow) and mitochondrial (HSP60; magenta) markers. Location of mitochondrial targeting sequence (MTS) and peroxisomal targeting sequence (PTS) on the full-length isoforms of ACAD10 and ACAD11 are illustrated at the top. **(b)** Diagram of experimental workflow to test 4-HA metabolism in Hepa1-6 cells. **(c) 4**-phosphovaleryl-CoA levels detected in Hepa1-6 cells treated with 0 mM or 10 mM unlabeled 4-HV. **(d)** 4-phospho-DHA-CoA levels detected in Hepa1-6 cells treated with unlabeled 100 µM 4-HDHA conjugated to 1% fatty acid-free BSA (w/v). Results are represented as the mean signal intensity normalized to total µg protein per sample -/+ SD (n = 3 replicate experiments). **(e)** Model of 4-HA catabolism at an organellar level.

To clarify their subcellular localizations, we ectopically expressed human ACAD10:FLAG and FLAG:ACAD11 in COS7 and U2OS cells and imaged their distribution. Our imaging shows distinct localization of ACAD10/11 to separate organelles in both cell lines: ACAD10:FLAG localizes to mitochondria and FLAG:ACAD11 localizes to peroxisomes (Fig. 3a; Extended Data Fig. 5a). Localization of these paralogs to mitochondria and peroxisomes was disrupted when the positions of their FLAG tags were swapped to the opposite termini (Extended Data Fig. 5b).

In metazoans, FAO is conducted in both mitochondria and peroxisomes depending on the length of the fatty acid^38^. Thus, we hypothesized ACAD10/11 are tasked with catabolizing 4-HA species of different lengths according to their organellar localization. Canonically, mitochondria catabolize short-to long-chain fatty acids (approximately 4 – 18 carbons) and peroxisomes catabolize very long-chain fatty acids (>20 carbons). Using our Hepa1-6 CRISPR/Cas9 KO cell models, we evaluated the necessity of either ACAD10 or ACAD11 in the catabolism of two different 4-HAs, 4-HV (5-carbons long) and 4-HDHA (22-carbons long), by monitoring the formation of their respective 4-phosphoacyl-CoA intermediates (Fig. 3b). All cell lines supplemented with the 4-HAs produced the corresponding 4-hydroxyacyl-CoA intermediates (Extended Data Fig. 5c-d). When given 4-HV, cells specifically lacking *Acad10* were unable to generate 4-PV-CoA, whereas cells lacking *Acad11* produced 4-PV-CoA at levels comparable to wild type (Fig. 3c). Reciprocally, when supplemented with 4-HDHA, cells lacking *Acad11* failed to produce detectable 4-phospho-DHA-CoA (Fig. 3d), whereas cells lacking *Acad10* performed like wild type. These results suggest that ACAD10 functions in mitochondrial short-chain 4-HA catabolism and ACAD11 in peroxisomal very long-chain 4-HA catabolism (Fig. 3e).

### *Acad11*^-/-^ mice exhibit perturbed 4-HA metabolism

Previously, *Acad10* knockout mice were shown to develop a range of metabolic complications, including the accumulation of excess abdominal adipose tissue and fasting rhabdomyolysis^39^; however, the physiological consequences of ACAD11-deficiency are unknown. Here, we established a whole-body knockout of *Acad11* in the C57BL/6 mouse background to extend these analyses and directly determine whether ACAD11 mediates the catabolism of 4-HAs *in vivo* (Extended Data Fig. 6a,b). Mice were born at normal Mendelian frequency and did not display differences in total body mass according to genotype (Extended Data Fig. 6c,d).

FAO enzymatic deficiencies often result in the systemic accumulation of unprocessed FAO intermediates^40^. Therefore, we hypothesized whole body deletion of *Acad11* would lead to an endogenous elevation of 4-HAs that fail to enter the canonical FAO pathway. To test this hypothesis, we developed and applied a targeted LC-MS method to measure various hydroxylated free fatty acids in mouse plasma (Fig. 4a). Long-chain 4-HA species, notably 4-OH C10 and 4-OH C12, were significantly elevated in the plasma of male and female *Acad11* KO mice when fed either a standard chow diet or short-term ketogenic diet (Fig. 4b,c; Extended Data Fig. 6e,f). We also observe the accumulation of shorter 4-HA species, which we speculate to be catabolic intermediates of hydroxylated fatty acid precursors^12^. We find 4-HAs are specifically perturbed in *Acad11* KO mice and do not reflect a general elevation of hydroxylated fatty acids. For example, on standard chow, 4-OH C10 and 4-OH C12 exhibited the strongest accumulation in KO mice, whereas their corresponding 3-OH and 5-OH isomers were unperturbed across all genotypes in both sexes (Fig. 4d,e).

**Fig 4.**
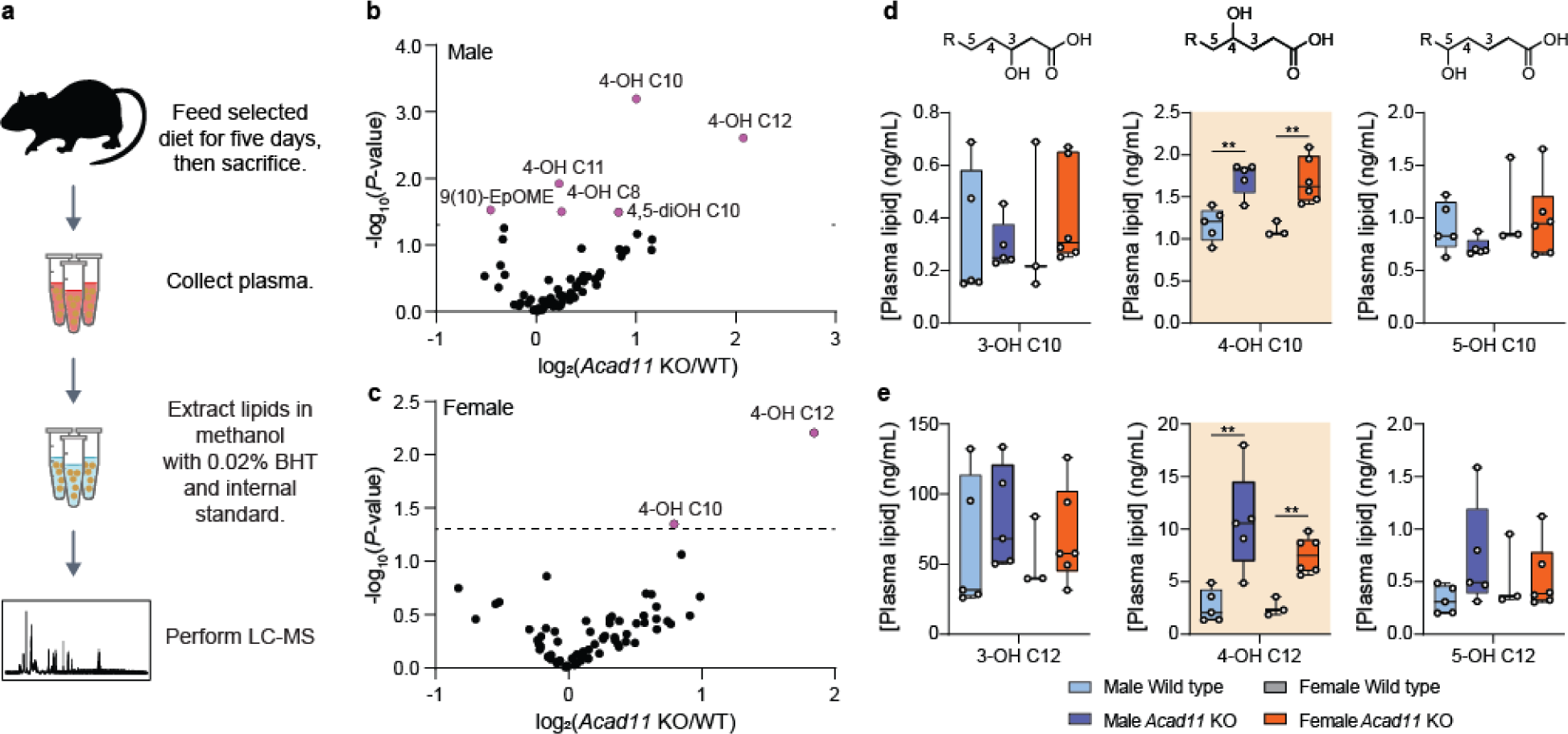
*Acad11*^-/-^ mice exhibit perturbed 4-HA metabolism. **(a)** Experimental design for quantitative measure of 4-HA abundance *in vivo.* **(b-c)** Volcano plots depicting log_2_-transformed fold changes of 4-HAs and other targeted hydroxylated lipids relative to statistical significance. Male and female KO mice were compared to respective littermate controls. Mice were fed a standard chow diet. **(d-e)** Plasma 4-HA concentrations of 4-OH C10 (top) and 4-OH 12 (bottom) across all genotypes and sex. 4-HAs are compared side-by-side with similar isomers.

Based on our animal model results, we propose that ACAD11 functions to eliminate 4-HAs *in vivo*. More broadly, our overall study nominates ACAD10 and ACAD11 as the primary gatekeepers of mammalian 4-HA metabolism, thereby providing a foundation to further investigate the mechanisms by which they catabolize 4-HAs, identify new 4-HAs connected to their activities, and explore the pathophysiology that may result from their disruption.

## Discussion

Here, we demonstrate that ACAD10 and ACAD11 catabolize 4-HAs through the major pathway described in literature^18,19^. ACAD10/11 are evolutionarily divergent members of the ACAD family^24^, possessing both an N-terminal kinase domain and unconventional hydrophilic residues in the ACAD active site. These features allow them to phosphorylate and process 4-hydroxylated acyl-CoA substrates, ultimately removing the hydroxyl group and producing 2-enoyl-CoAs that can enter the FAO cycle. The presence of hydrophilic residues is uncommon in ACAD active sites, which are typically hydrophobic to accommodate neutral-charged, long-chain acyl tails^32^. However, our structure-guided analysis of ACAD11 reveals an HCMR motif that is essential for activity and exists proximal to FAD and the catalytic aspartate. We also identify a highly conserved arginine upstream of the HCMR motif (R626 for mouse ACAD11) that is likely required for stabilizing the negatively charged phosphate during its release. Interestingly, the decarboxylation activity of glutaryl-CoA dehydrogenase — another atypical member of the ACAD family — relies on an analogous arginine, which is thought to stabilize the negatively charged carboxylate group^41^.

The precise catalytic mechanisms of both ACAD10/11 domains remain unclear. In ACAD11’s ACAD domain, FAD is bound in the canonical position seen in other ACADs and is required for catalytic turnover of 4-phosphoacyl-CoA. Although 4-phosphorylated substrates and their corresponding 2-enoyl-CoAs products have the same oxidation state, our experiments with deazaFAD suggest FAD still acts in a redox active manner. However, FAD is not terminally reduced to FADH_2_, as it does not require an electron acceptor to be regenerated post-catalysis *in vitro*. We suggest a mechanism by which FAD is first reduced, but later donates a hydride as part of the phosphoelimination to return to an oxidized state. Similar “rebound” functionality for flavin cofactors has been observed in other enzymatic mechanisms, including those of chorismate synthase and a linoleic acid isomerase from *Propionibacterium acnes*^42,43^. Further work is required to clarify the mechanistic roles of FAD and the other unique active site features of ACAD10/11 revealed through our structure.

ACAD10/11 are localized to mitochondria and peroxisomes, respectively. Although catalytically similar, our work in cultured cells and *in vivo* suggests that ACAD10/11 do not fully compensate for one another in the catabolism of 4-HAs. We propose that this subcellular partitioning broadens the range of 4-HA species that the cell can catabolize. 4-HAs detected thus far in human and rodent models range from 4 to 22 carbons^12,14,15,17^; conventional catabolism of fatty acids spanning this range requires enzymes from both mitochondria and peroxisomes^38^. Additionally, ACAD10/11 may function with co-localized FAO machinery to catabolize precursor fatty acids that are eventually shortened into 4-HAs, such as those with hydroxyl group on even-numbered carbons^21^. For example, microorganisms that are fed ricinoleic acid (i.e., 12-hydroxyoleic acid) produce 4-OH C10 as a catabolic intermediate that is further degraded^16^.

But why do cells need to catabolize 4-HAs? We speculate that ACAD10- and ACAD11-dependent catabolism of 4-HAs may be relevant for the efficient clearance of oxygenated lipids, such as those produced during excessive oxidative stress or inflammation. The origins of 4-HAs are largely unknown, however a few species may be produced from reactive aldehydes, such as 4-hydroxynonenal (4-HNE), that form during lipid peroxidation^44^. Previous work with rat liver tissue identified a nine-carbon 4-phosphoacyl-CoA intermediate derived from 4-OH C9, a downstream intermediate of 4-HNE^18,21^; thus, FAO has been proposed as a pathway for eliminating 4-HNE and protecting against its cytotoxic effects. In mammalian cells, mitochondrial and peroxisomal FAO coordinate the catabolism of oxylipins^45^, which are signaling molecules that control inflammation and vascular response^46^. For some (e.g., 12-HETE), their catabolism may yield 4-HA intermediate bottlenecks that are managed by ACAD10/11^47^.

To date, no monogenic disorders stemming from either ACAD10 or ACAD11 have been identified. However, several genome-wide association studies have identified single-nucleotide polymorphisms in both paralogs associated with coronary heart disease, obesity, diabetes, and insulin resistance in diverse human populations^48-51^. Our identification of ACAD10/11 as the primary gatekeepers of 4-HA metabolism will empower further studies of 4-HA function and metabolism to reveal their physiological relevance to human health and disease.

## Methods

### Animal models

Cryopreserved *Acad11*^-/-^ (C57BL/6NJ) mouse sperm was purchased from The Jackson Laboratory (51098-JAX). *In vitro* fertilization of C57BL/B6J donor eggs with the sperm was performed by the University of Wisconsin-Madison Animal Models Core. *Acad11* heterozygous progeny were inbred to maintain colony. Wild-type littermate controls were included in all experiments. Breeder mice were maintained on a high energy diet (Formulab Diet 5015). Experimental mice were maintained on standard chow diet (Formulab Diet 5008) unless specified otherwise. Mice were housed under a standard 12-hour dark/12-hour light cycle and had free access to water and food. Our animal protocol was approved by the Institutional Animal Care and Use Committee of the College of Agricultural and Life Science at the University of Wisconsin-Madison.

### Mouse genotyping

Mouse tail clippings were digested (24 – 48 hrs, 55°C) with 10 µL of 10 mg/mL proteinase K in 400 µL of 20 mM Tris-HCl, 25 mM EDTA, 100 mM NaCl, 0.5% SDS (v/v), pH 8.0. The next day, samples were centrifuged to pellet undigested matter (16300 x *g*, 5 mins). 390 µL of supernatant was mixed with 800 µL of cold isopropanol by inverting tubes twenty times.

Samples were centrifuged to pellet precipitated genomic DNA (16300 x *g*, 10 mins, 4°C). After aspirating the supernatants, precipitates were washed with 1 mL of 100% ethanol followed by 1 mL 70% ethanol. After drying in a fume hood, genomic DNA precipitates were resuspended in 1X TE buffer, pH 8.0. *Acad11* genotypes were assessed using a three-primer PCR strategy with the 5X Multiplex PCR Mix (NEB) to allow for the simultaneous detection of both wild-type and mutant alleles. PCRs were amplified by the following steps (steps 2-4 were repeated 35 times): step 1) initial denaturation (1 min, 95°C); step 2) denaturation (30 sec, 95°C); step 3) annealing (30 sec, 53°C); step 4) elongation (1.5 min, 68°C); and step 5) final extension (5 min, 68°C). PCRs were resolved on a 2% agarose gel stained with EtBr prior to imaging.

### Cell culture

Hepa1-6 cells (mouse hepatoma; ATCC; CRL-1830) were cultured in high glucose DMEM containing 10% fetal bovine serum (FBS) (Gibco) and maintained at 37°C and 5% CO_2_. Cells used for experiments underwent no more than six passages. Monoclonal CRISPR knockout Hepa1-6 cell lines were generated by the Genome Engineering and Stem Cell Center (GESC) at Washington University in St. Louis using two different single-stranded guide RNAs that targeted early exons of *Acad10* and *Acad11*. After selection of cells with indels predicted to introduce early stop codons and verification by next generation sequencing, two monoclonal lines per gene knockout combination were chosen. Double knockout (DKO) cells were generated from *Acad11* KO-b. Cell lines were verified to be mycoplasma-free with the LookOut Mycoplasma qPCR detection kit (Sigma-Aldrich).

### Multiple sequence alignment and homology searches

Computational search for *lva* operon enzyme homologs in mammals was conducted by querying their primary amino acid sequences against reference proteomes available on UniProtKB/SwissProt databases (taxonomy restricted to “mammalia”) using the BLASTp 2.12.0+ algorithm. Subsequent analyses of LvaA and LvaC sequence conservation and coverage were performed with primary sequence sets containing NCBI HomoloGene sequences of ACAD10/11 or the full-length sequences of human acyl-CoA dehydrogenases. Multiple sequence alignments of FASTA formatted sequences were generated using the default settings of the MAFFT algorithm (Version 7.505)^52^. The MAFFT alignment results were visualized with Jalview (Version 2.11.2.5). The Pfam database was referenced to illustrate the human enzyme domain architecture in Figure 1b.

### Plasmid cloning

For recombinant expression vectors, the coding sequences of mouse *Acad10* and *Acad11* were amplified from wild-type female C57BL/6J mouse liver tissue cDNA by PCR. Using Gibson assembly, *Acad10^NΔ34^*(gene product lacking 34 amino acid N-terminal mitochondrial targeting sequence) and *Acad11* were cloned into different expression vectors. *P. pastoris* codon-optimized *Acad10^NΔ34^* was ordered as gene fragments from GenScript and incorporated into the pPicZb vector with a 3’ GFP tag coding sequence and *Acad11* was cloned into the pE-SUMOstar vector with a 5’ SUMO tag coding sequence. pE-SUMO vectors for recombinant mouse ACADM^NΔ23^, ACADS^NΔ29^, and ACADVL^NΔ41^ expression and purification were cloned similarly from templates purchased from GenScript. For ectopic gene expression in cell culture, human *ACAD10* and *ACAD11* sequences were cloned into pcDNA3.1 vectors with N-terminal or C-terminal FLAG sequences using similar strategies. Genes were amplified by PCR and inserted into the pcDNA3.1 backbone using Gibson assembly following digestion with EcoRI and HindIII. For all cloning work, the 2X Q5 Master Mix (NEB) was used for high-fidelity PCR and the 2X HiFi Gibson Assembly Mix (NEB) was used for amplicon assembly. Point mutations were generated using the Q5 Site-Directed Mutagenesis Kit (NEB). Primer sequences and plasmid construct information is listed in Supplementary Table 1.

### Bacterial recombinant protein purification

For purification of *M. musculus* ACAD11, ACADM^NΔ23^, ACADS^NΔ29^, and ACADVL^NΔ41^, pE-SUMO constructs were transformed into B21-CodonPlus (DE3)-RIPL competent *E. coli*. Colonies were picked and cultured in LB containing 50 μg/mL kanamycin and 25 μg/mL chloramphenicol and expression was induced with 100 µM IPTG (overnight with shaking, 18°C). Cultures were harvested via centrifugation (4000 x *g*, 20 min, 4°C) and pellets were resuspended in lysis buffer (50 mM HEPES, 300 mM NaCl, 5% glycerol, 1 mM MgCl_2_, 500 μM phenylmethylsulfonyl fluoride (PMSF), 1 mM β-mercaptoethanol (β-ME), pH 7.4). For purification of ACAD11 constructs and ACADVL^NΔ41^, 0.5% NP-40 IGEPAL (Sigma-Aldrich) and 0.5% Tween-20 (Fisher Scientific) were included in the lysis buffers respectively. Cell pellets were suspended by hand into a slurry and sonicated on ice with a Bronson sonifier (75% amplitude, 20 sec on and 5 min rest, 3 cycles). Lysates were clarified by a hard spin (30000 x *g*, 30 min, 4°C). The soluble fractions were incubated with 3-10 mL of Ni-NTA resin in buffer without detergent for 1 hr with gentle tube rotation at 4°C. Supernatants were removed and the resins were washed with buffer containing increasing concentrations of imidazole (10 mM – 50 mM), followed by elution with 25 mL of buffer containing 300 mM imidazole. The eluates were dialyzed overnight in the presence of Ulp1 protease (1 U/mg protein). The next day, the dialyzed protein samples were incubated with 1 mL of Ni-NTA resin (1 hr, 4°C) to remove excess free protease, SUMO tag, and contaminants. The supernatants were collected and concentrated to 1 mL using an Amicon centrifugal filter with a 50 kDa cut-off. Concentrated proteins were separated by size exclusion chromatography, and fractions of the first major elution peak were assayed for the presence of recombinant protein via SDS-PAGE according to standard procedures. The purest fractions were pooled and exchanged into storage buffer (50 mM HEPES, 300 mM NaCl, 20% glycerol, 1 mM MgCl_2_, 1 mM β-ME, pH 7.4) before flash-freezing with liquid nitrogen and storing at −80°C. The expression and batch purification of recombinant His-LvaE was performed as previously described except with the buffers used to purify recombinant ACAD11^22^. Protein concentration was quantified via Bradford assay (Bio-Rad).

### Yeast recombinant protein purification

For purification of *M. musculus* ACAD10^NΔ34^, pPicZ-*Acad10^NΔ34^*-GFP was transformed and expressed in *Pichia pastoris*. Individual clones were selected, and expression of the GFP-tagged protein was verified by western blot using an anti-GFP antibody (Sigma-Aldrich). Colonies with the highest expression for a given construct were used to inoculate 300 mL of YEPD media containing 200 µg/mL Zeocin and grown overnight with shaking (220 rpm, 30°C). The next day, the full culture volume was used to inoculate 12 L of YEPD containing 200 µg/mL Zeocin and grown for 24 hours with shaking (220 rpm, 30°C). Turbid cultures were centrifuged (20 min, 4000 x *g*, RT) and the pellet was gently resuspended in buffer minimal methanol medium (BMM) (200 mM potassium phosphate, 1X yeast nitrogen base, 40 µg biotin, 0.7% methanol, 100 µg/mL zeocin, pH 6.0) to induce ACAD10^NΔ34^-GFP-His protein expression (220 rpm, 24 hrs, 24°C). Cultures were harvested via centrifugation (4000 x *g*, 20 min, 4°C) and extruded from a wide-bore syringe into liquid nitrogen to create frozen droplets. Droplets were subjected to lysis via CryoMill under frozen conditions. CryoMill lysate powder was gently resuspended in chilled lysis buffer and clarified by a hard spin (30000 x *g*, 30 min, 4°C). The supernatant was collected and incubated with 10 mL Ni-NTA resin in standard purification buffer with gentle tube rotation (1 hr, 4°C). After removing supernatant, the resin was washed with increasing concentrations of imidazole (10 mM – 50 mM) followed by elution with buffer containing 300 mM imidazole. The eluate was dialyzed overnight with PreScission protease added (1 U/mg protein). The following day, the dialyzed protein was carefully stirred and diluted 20-fold in standard protein buffer without NaCl, resulting in a final NaCl concentration of 15 mM. The diluted mixture was concentrated to 25 mL using an Amicon centrifugal filter and run continuously (0.5 mL/min) over a 5 mL HiTrap® SP Fast Flow FPLC column (Millipore Sigma) for 2-3 hours to maximize protein capture. The protein was eluted from the column using a gradient of low to high salt buffer and SDS-PAGE was used to identify fractions containing the protein of interest, which were then pooled. The pooled fractions were stored overnight at 4°C on ice before being concentrated to 1 mL using an Amicon centrifugal filter with a 50 kDa cut-off. Like the purification of recombinant proteins from bacteria, the concentrated protein was separated by size exclusion chromatography. After verifying the presence of protein by SDS-PAGE, fractions of the main elution peak were pooled and exchanged into storage buffer prior to flash-freezing and storage at −80°C.

### Production of acyl-CoA enzyme substrates

4-hydroxy acids (4-HAs) used in this study were generated from ɣ-lactone precursors (Supplementary Table 3), unless otherwise specified. Lactones were saponified in 20-30% molar excess NaOH at 60°C – 75°C with intermittent vortexing. For *in vitro* experiments, short-chain acyl-CoAs were generated *in situ* by incubating 0.5 µM recombinant LvaE with equimolar amounts of ATP, CoA, and free fatty acid (i.e., valerate, 4-hydroxyvalerate, 2-pentenoate, or 3-pentenoate) for 30 mins at room temperature (50 µL volume). Reactions containing acyl-CoA products were used directly as substrates for *in vitro* activity assays and standards for HPLC method development.

### *In vitro* enzyme activity assays

For LC-MS analysis, 100 µL reactions were prepared with 50 μM ATP, 50 μM FAD, 50 μM acyl-CoA substrate, and 0.5 μM recombinant protein. Reactions were then rotated at 37°C for 30 minutes. Reactions were quenched with 100 μL of cold quenching solvent (MeOH:Water (1:1, v/v) containing 5% acetic acid). Samples were centrifuged to pellet any precipitate (16300 x *g*, 10 mins, 4°C). Each supernatant was transferred to a new 1.5 mL tube and dried in a speed vacuum concentrator overnight. Samples were stored at −80°C prior to reconstitution in 500 µL LC-MS grade water before analysis. For HPLC analysis, a similar protocol was followed with minor changes. Reactions were assembled with 500 μM ATP, 5 μM FAD, 50 μM acyl-CoA substrate, and 0.5 μM recombinant protein unless otherwise noted. The final assay volume was 25 μL and assays were quenched with 25 μL quenching solvent. Quenched reaction mixtures were transferred directly to amber glass vials with inserts. HPLC elution standards were purchased or synthesized with enzymes and diluted in quenching solvent. All conditions were measured in triplicate and tested alongside negative control reactions lacking recombinant enzyme.

### High performance liquid chromatography (HPLC) analysis of *in vitro* assay mixtures

HPLC analysis of acyl-CoA species was performed similarly to previously published methods^53^. Prior to sample runs, column (Betasil C18, 100 × 2.1 mm, 3 µm particle; Thermo Scientific) was washed with 100 mL of Solvent B (MeOH) followed by 100 mL of Solvent A (20 mM ammonium acetate:MeOH (94:6, v/v), pH 7.0). A methanol “blank” was injected at the start of each running period. The injection volume for all samples, blanks and standards was 15 μL. The flow rate for the method was 0.5 mL/min and the total duration of each individual run was 1 hr. The solvent percentages were as follows for Solvent A: 10.5 min of a gradient from 100% to 94%, 7.5 min at 94% isocratic, 27 min of a gradient from 94% to 25%, 2 min of a gradient from 25% to 100%, and 13 min of 100% isocratic. Acyl-CoA peaks were quantified by post-column UV-Vis absorbance measurements at 259 nm. Peak identities were verified using corresponding LvaE-generated standards as described above and abundance was quantified using the Chromeleon 7.2.10 software with Cobra Wizard defaults. Synthetic standards of 4-phosphoacyl-CoAs could not be synthesized, thus their elution times were confirmed by injecting reaction mixtures containing LvaE-generated 4-hydroxyacyl-CoAs and ACAD-inactive recombinant ACAD11 (A*). The presence of 4-phosphoacyl-CoA in these reactions was confirmed by LC-MS.

### Assay with free fatty acid substrates using ADP-Glo™

The ADP-Glo™ assay (Promega) was performed according to the manufacturer’s instructions with the following modifications. Free fatty acid substrates were saponified at 2 M in 20% molar excess NaOH and diluted to 20 mM before use. All solutions were diluted in assay buffer (50 mM HEPES, 1 mM MgCl_2_, 1 mM DTT, pH 7.4). In 96-well plates, 0.5 µM recombinant protein was mixed with 500 µM ATP, 5 µM FAD, and 500 µM free fatty acid or LvaE-generated acyl-CoA substrates. Reactions were incubated at 37°C for 30 min. ADP-Glo Reagent (5 μL) was added and incubated with a cover slip for 40 min at room temperature. Kinase Detection Reagent was then added (10 μL) and incubated for 60 min at room temperature. Luminescence was read using default values on a Biotek Cytation 3 plate reader. An ADP/ATP standard curve was made according to the manufacturer’s instructions using reagents supplied in the kit. All conditions were measured in triplicate.

### Acyl-CoA dehydrogenase ferrocenium assay

Ferrocenium assays were conducted as previously described with some modifications^25^. Ferrocenium hexafluorophosphate (Sigma-Aldrich) was dissolved in 10 mM HCl and quantified using Nanodrop pedestal UV-Vis spectroscopy. Substrate stock solutions (hexanoyl-CoA (C6:0), decanoyl-CoA (C10:0), myristoyl-CoA (C14:0), stearoyl-CoA (C18:0), and behenoyl-CoA (C22:0) (Avanti Polar Lipids)) were dissolved in assay buffer (50 mM HEPES, 1 mM MgCl_2_, 1 mM DTT, pH 7.5). Reaction mixtures in assay buffer were prepared in a 96-well plate with 250 µM ferrocenium, 0.5 µM FAD, and 250 µM substrate. Each reaction was initiated by adding enzymes (ACAD11 wild type, mutant, ACADM, ACADS, ACADVL) at a final concentration of 0.5 μM. The decrease in ferrocenium absorbance at 300 nm was measured as a function of time for five mins using the maximum speed sweep read setting in a Biotek Cytation 3 plate reader. Activity was calculated in units of ΔAbs/min based on the slope of the linear portion of the enzyme kinetic curves, as determined in Microsoft Excel and GraphPad Prism. All conditions were measured in quadruplicate.

### Cryo-electron microscopy (EM) sample preparation

For cryo-EM, recombinant ACAD11 (K* and A* mutant) was purified from bacteria as described previously with the following modifications. The supernatant from the second Ni-NTA resin incubation step was collected and diluted 20-fold in standard protein buffer without NaCl, resulting in a final NaCl concentration of 15 mM. The diluted solution was subjected to cation exchange chromatography as described for recombinant ACAD10 above. The eluate from this step was collected, pooled, and subjected to size exclusion chromatography as described previously with the following changes. The size exclusion chromatography buffer used was 50 mM HEPES, 50 mM NaCl, 2% glycerol, 50 μM FAD, pH 7.5. The final sample was also amended to include 100 μM 4-hydroxyvaleryl-CoA for the kinase-dead mutant or 4-phosphovaleryl-CoA for the ACAD-dead mutant, generated using LvaE and ACAD-dead ACAD11 as described previously. The purest 1.5 mL fraction of eluate was collected and concentrated in an Amicon centrifugal filter before freezing. The final protein concentrations were 230 µg/mL for the kinase-dead mutant and 388 μg/mL for the ACAD-dead mutant. Cryo-EM samples were prepared on Quantifoil copper 300-mesh 2/2 holey carbon grids and were plunge-frozen in a Vitrobot Mark IV (Thermo Fisher Scientific) with a chamber temperature of 4°C and a humidity set to 95%. Prior to freezing, grids were freshly plasma cleaned on a Gatan Solarus 950 plasma cleaner for 60s. 3 µL of sample was adsorbed onto the grids for 20s, then blotted for 2s and immediately plunge-frozen in liquid ethane.

### Cryo-EM data processing and map generation

For ACAD11 K* mutant and A* mutant datasets, 2,789 and 2,965 raw movies respectively were imported to cryoSPARC V3.3.1^54^ and subjected to patch motion correction and contrast transfer function (CTF) estimation. The high-quality images were manually selected for subsequent data processing. Particles were initially picked using blob picker. Subsequent 2D classes were used as templates for template picker. 722,751 particles of ACAD11 K* mutant and 1,294,186 particles of ACAD11 A* mutant were extracted with a box size of 300 pixels. Classes containing high-resolution particles were selected and utilized to generate initial models. For the ACAD11 K* mutant, 242,249 final particles were subjected to non-uniform refinement with imposed D2 symmetry. For the ACAD11 A* mutant, heterogenous refinement was conducted to further remove low-quality particles. 297,619 final particles were subjected to local refinement with imposed D2 symmetry. The subsequent two half maps were imported to deepEMhancer 0.14^55^. The maps were post-processed at highRes mode to further improve quality for model building.

### Cryo-EM model building and refinement

For the ACAD11 K* mutant, the crystal structure of human ACAD11 (PDB:2WBI), and the AlphaFold^56^ model for ACAD11 were docked into the cryo-EM map and adjusted with rigid-body fit and real-space refinement in COOT 0.9.8^57^. The model was further refined in PHENIX 1.20.1^58^. For the ACAD11 A* mutant, the ACAD11 K* mutant model and the AlphaFold model were docked into the map, followed by rigid body fit in COOT 0.9.8 and real-space refinement in PHENIX 1.20.1. Final models were validated using MolProbity^59^ and figures were generated with ChimeraX 1.5^60^. The 3D structure of octanoyl-CoA (identifier CO8) was obtained from PDBeChem^61^ and used for *ab initio* surface-based docking in HADDOCK 2.4^31^. The isolated “C” chain of the ACAD11 A* structure was used as the protein target and default parameters were maintained, with the addition of the “Random patches define randomly ambiguous interaction restraints from accessible residues.” The complete HADDOCK 2.4 report can be found in Supplementary Figure 1.

### Conjugation of 4-HDHA to fatty acid-free BSA

20 mM 4-HDHA (synthesized by WuXi) in ethanol was dried under nitrogen gas for ∼1 hr. The dried residues were then reconstituted in high glucose DMEM containing 10% fatty acid-free BSA (w/v). Samples were sonicated for 5 mins in a 37°C water bath using the degas setting, followed by 5 mins of rest (four cycles total, inverting tubes between each cycle). To prepare working stocks, conjugated 4-HDHA was diluted 10-fold in high glucose DMEM containing no BSA and sterilized with a 0.22 µM filter. Media was always prepared fresh and used the same day.

### 4-HA catabolism studies in Hepa1-6 cells

Hepa1-6 cells were trypsinized at confluency and counted using the CytoSMART automated cell counter. One million cells were seeded in individual 10 cm plates (two plates prepared per sample condition to increase sample amount). After 48 hours, cells were rinsed with 1X dPBS (Gibco) and supplemented with high glucose DMEM containing 10% dialyzed FBS (Bio-Techne; S12850H). For studying 4-hydroxyvalerate (4-HV) metabolism, cells were given fresh medium containing 5 mM [^13^C_5_]-4-HV (synthesized by WuXi) or 10 mM unlabeled 4-HV sodium salt (Santa Cruz Biotechnology). Cells were cultured with 4-HV for 6 hours prior to harvest. For studying 4-hydroxydocosahexaenoate (4-HDHA) metabolism, cells were supplemented with high glucose DMEM containing 100 µM 4-HDHA conjugated to 1% fatty acid-free BSA (w/v) (Sigma-Aldrich; A6003). Cells were cultured with 4-HDHA for 24 hours prior to harvest. All conditions were tested as three independent experiments alongside negative controls treated with no exogenous fatty acid.

### Solid-phase extraction of 4-phosphoacyl-CoA and other acyl-CoAs

Acyl-CoAs were extracted from Hepa1-6 cells using adapted solid-phase extraction (SPE) procedures^18,62^. For studies with 4-HV, cells were rinsed twice with cold 1X dPBS and then were scraped into 1 mL of cold extraction solvent (MeOH:H_2_O (1:1, v/v) with 5% glacial acetic acid). After transferring to a 1.5 mL screw-cap microcentrifuge tube, cells were vortexed for 20 seconds and pelleted (16300 x *g*, 10 mins, 4°C). While spinning, 100 mg 2-(2-pyridyl)-ethyl silica gel cartridges (Supelco) were activated with 1 mL methanol followed by 1 mL extraction solvent. 2 mL of total clarified cellular extract from two identical 10 cm plates was passed through the cartridge. The SPE matrix was washed once with 1 mL MeOH:50 mM ammonium formate (pH 6.3) (1:1, v/v). Compounds were eluted with 0.5 mL MeOH:50 mM ammonium formate (pH 6.3) (4:1, v/v) twice and 0.5 mL MeOH twice. 2 mL of the total eluate was dried in a speed vacuum concentrator. Dried extracts were stored at −80°C. Prior to LC-MS analysis, Hepa1-6 cell acyl-CoA extracts were reconstituted in 50 µL of water:MeOH (97:3, v/v) with 10 mM tributylamine (pH 8.2; adjusted with 10 mM acetic acid).

For studies with 4-HDHA, cells were rinsed twice with cold 1X dPBS, scraped into 0.9 mL of cold acetonitrile/IPA (3:1, v/v), and transferred to 2 mL screw-cap tubes. Samples were vortexed for 30 seconds. 0.3 mL of 0.1 M KH_2_PO_4_, pH 6.7 was added to each sample followed by vortexing for 30 seconds. The samples were then centrifuged (16300 x *g*, 10 mins, 4°C). 1.2 mL of the supernatants were transferred to new 2 mL screw-cap tubes and were acidified with 300 µL of glacial acetic acid followed by vortexing for 15 seconds. SPE cartridges were pre-conditioned with 1 mL MeOH followed by 1 mL MeCN/IPA/water/acetic acid (9:3:4:4, v/v/v/v). 1.2 mL of each acidified sample was loaded onto the cartridge (n = 2 technical replicates were loaded sequentially). The cartridges were washed once with 1.2 mL of MeCN/IPA/water/acetic acid (9:3:4:4, v/v/v/v). Acyl-CoAs were eluted with 1 mL MeOH/250 mM ammonium formate (4:1, v/v) three times. Only the second two elutions were collected based on the elution profile of a previous experiment. 2 mL of eluate was transferred to a new 2 mL screw-cap tube and dried by speed vacuum. Dried residues were immediately stored at −80°C. Prior to LC-MS analysis, dried extracts were resuspended in 50 µL of 50 mM ammonium acetate, 20% MeCN. For data normalization, the residual pellets were dried, boiled in 500 µL of 0.2 M NaOH at 95°C, and assayed for total protein mass with the Pierce BCA Protein Assay Kit (Thermo Scientific).

### Mouse plasma collection and 4-HA extraction

Male and female mice (13-15 weeks old) were fed a standard chow (Envigo; TD.00606) or ketogenic diet (Envigo; TD.96355) for five days. After fasting for 6 hrs with access to water, mice were anesthetized under isoflurane and euthanized by decapitation. Blood was immediately collected into tubes containing sodium citrate buffer (Covidien). Plasma was isolated by centrifugation (10 mins, 2000 x *g*, 4°C) and the supernatants were transferred to fresh tubes. Plasma samples were flash-frozen in liquid nitrogen and stored at −80°C until day of extraction. Methanol precipitation was used to extract plasma 4-HAs. After thawing on ice, 50 µL of plasma was mixed with 1 mL of cold MeOH spiked with internal standards. After vortexing for 30 seconds, samples were placed at −80°C for 1 hr. Samples were centrifuged to pellet insoluble material (16300 x *g*, 10 mins, 4°C). 950 µL of each supernatant was transferred to a new 1.5 mL tube and dried overnight in a speed vacuum concentrator. Dried samples were reconstituted in 50 µL of MeOH:Water (1:1, v/v) prior to LC-MS analysis.

### LC-MS of acyl-CoA metabolomics and stable isotope tracing

For *in vitro* enzyme activity and cell culture experiments, LC-MS analyses of 4-HV acyl-CoA intermediates were conducted using a Vanquish ultra-high-performance liquid chromatography (UHPLC) system (Thermo Scientific) coupled to a hybrid quadrupole-Orbitrap™ mass spectrometer (Thermo Scientific; Q Exactive™) equipped with electrospray ionization operating in negative-ion mode. The chromatography was performed at 25°C using a 2.1 × 100 mm reverse-phase C_18_ column with a 1.7 μm particle size (Water™; Acquity UHPLC BEH). The chromatography gradient used Solvent A (97:3 H_2_O:methanol with 10 mM tributylamine adjusted to pH 8.2 using 10 mM acetic acid) and Solvent B (100% methanol) as follows: 0-2.5 mins, 5% B; 2.5-5 mins, linear gradient from 5% B to 20% B; 5–7.5 mins, 20% B; 7.5–13 mins, linear gradient from 20% B to 55% B; 13–15.5 mins, linear gradient from 55%–95% B; 15.5-18.5 mins, 95% B, 18.5-19 mins, linear gradient from 95% B to 5% B; 19–25 mins, 5% B. The flow rate was held constant at 0.2 mL/min. For the metabolomics method, Full MS-SIM (single ion monitoring) and Parallel Reaction Monitoring (PRM) methods were run with an inclusions list. Eluent from the column was injected into the MS for analysis from 2 min to 17.2 min, at which point flow was redirected to waste for the remainder of the run. Properties for the MS methods included the following: Scan range between 500 and 1,000 *m*/*z*; automatic control gain (ACG) target 5e6; maximum injection time (IT) 100 ms; a resolution of 70000 full width at half maximum (FWHM); and an isolation window of 1.4 and 12 *m*/*z* for the PRM method for the non-isotopically labeled and isotopically labeled experiments, respectively. Fragmentation of CoA, pentenoyl-CoA, isovaleryl-CoA, 3-HV-CoA/4-HV-CoA, and 4-PV-CoA was achieved by using a normalized collision energy of 25 and a scanning window from 14.5–17 mins. The autosampler and column compartment were kept at 4°C and 30°C, respectively. Data analysis was performed using the El-MAVEN (Elucidata)^63^ and the Xcalibur™ (Thermo Fisher Scientific) softwares. Compounds were identified based on retention times matched to pure standards or via fragmentation patterns reported by Rand et al., 2017 (Supplementary Table 2).

LC-MS measurement of 4-HDHA acyl-CoA intermediates was performed using a Thermo Vanquish Horizon UHPLC system coupled to a Thermo Exploris 240 Orbitrap mass spectrometer. For LC separation, a Vanquish binary pump system (Thermo Scientific) was used with a Waters Acquity Premier CSH Phenyl-Hexyl column (100 mm × 2.1 mm, 1.7 µm particle size) held at 30°C under 300 µL/min flow rate. Mobile phase A consisted of MeCN/Water (5:95, v/v) with 10 mM Ammonium Acetate. Mobile phase B consisted of MeCN/Water (95:5, v/v). For each sample run, mobile phase B was held at 2% for the first 1.5 mins then increased to 15% over the next 1.5 mins. Mobile phase B was then further increased to 95% over the following 2.5 mins and held for 9 mins. The column was then re-equilibrated for 5 mins at 2% B before the next injection. 10 µL of sample was injected by a Vanquish Split Sampler HT autosampler (Thermo Scientific) while the autosampler temperature was kept at 4°C. The samples were ionized by a heated ESI source kept at a vaporizer temperature of 350°C. Sheath gas was set to 50 units, auxiliary gas to 8 units, sweep gas to 1 unit, and the spray voltage was set to 3500 V using negative mode. The inlet ion transfer tube temperature was kept at 325°C with 70% RF lens. The identity and retention time of the acyl-CoA derivatives and phosphorylated intermediate were first confirmed from cells fed high purity 4-HDHA using a parallel reaction monitoring (PRM) method with HCD at 30%, targeting selected ion fragments generated from the fragmentation of the hydrogen loss (H-) ion. Select fragments were chosen based on previous studies examining the phosphorylated intermediate of shorter-chain 4-hydroxyacyl-CoAs^22^. Quantification of experimental samples was performed using full scan mode at a resolution of 60000, targeting the hydrogen-loss ion of 4-hydroxy-DHA-CoA (m/z = 1092.3325) and 4-phospho-DHA-CoA (m/z = 1172.2988). Peak integration was performed using Tracefinder 5.1 (Thermo Scientific).

For all measurements, raw intensities of verified compound peaks were normalized to total protein mass of the sample and were used as a proxy for compound abundance in cells.

### LC-MS of mouse plasma lipids

Lipid extracts were analyzed using an Agilent Infinity II LC coupled to an Agilent 6495c triple quadrupole mass spectrometer as recently described^64^. The LC was equipped with an Agilent RRHD Eclipse Plus C18 column (2.1 × 150 mm, 1.8 µm) with an Agilent Eclipse Plus guard column (2.1 × 5 mm, 1.8 µm) kept at 50°C. Mobile phase A consisted of 0.1% acetic acid and mobile phase B was MeCN:IPA (90:10, v/v). The LC gradient was as follows: start at 15% B to 33% at 3.5 min, to 38% B at 5.5 min, to 42% B at 7 min, to 48% B at 9 min, to 65% B at 15 min, to 75% B at 17 min, to 85% B at 18.5 min, to 95% B at 19.5 min, and finally to 15% B at 21 min, held until 26 min at a constant flow of 0.350 mL/min. Multi-sampler was kept at 4°C and a blank injections were run between every sample. Samples were injected in random order and within 48 hrs of extraction.

MS analysis was in negative ionization with the following parameters: drying gas at 290°C at 10 L/min, nebulizer at 35 psi, sheath gas at 350°C at 11 L/min, capillary voltage at 3500 V and nozzle voltage at 1000 V. A dynamic multi-reaction monitoring (dMRM) method was used for analysis. A library of retention times and transitions was created using synthesized 4-HA standards and other oxygenated lipids from commercial mixes. Additional oxylipins for which standards were not tested were programmed using previously published transitions and retention time correlations^45,65^. Several collision energies between 10 – 40 V were tested to determine transitions for oxylipin standards that lack known transitions or were not detected using previously published transitions. Plasma pooled from all experimental mice that were spiked with all external standards was used to refine and finalize the method for plasma matrix. Additionally, external standards were injected preceding and following samples to ensure there were no retention time shifts or changes in detected analyte abundances due to instrument variability. Retention times and transitions used to identify and quantify targeted molecules, as well as the internal standards used for quantitation, are described in Supplementary Table 2. Data was collected and analyzed using the Agilent MassHunter Suite. Automated peak picking was used to integrate peaks with a signal-to-noise ratio > 3.0 and retention times within 0.30 min of the expected retention time. All peaks were then manually assessed for appropriate shape. Any samples with peak height less than background were removed. Compounds present in at least 70% of samples were reported in the final dataset in units of “ng lipid per mL plasma” after normalization to the appropriate internal standard.

### Microscopy

COS-7 (ATCC; CRL-1651) and U-2 OS (ATCC; HTB-96) cells were cultured in DMEM containing 10% FBS at 37°C and 5% CO_2_. Cells were seeded on 12 mm (1 ½ thickness) coverslips and incubated for 24 hrs prior to transfection. Cells were transfected with either human full-length ACAD10:FLAG, FLAG:ACAD10, ACAD11:FLAG, or FLAG:ACAD11 expressed under a CMV promoter in pcDNA3.1 constructs using Lipofectamine 3000 (Thermo Scientific) in OptiMEM (Gibco). Cells were cultured for 24 hrs prior to fixation with 4% paraformaldehyde in OptiMEM for 5 mins. Followed by washing with 1X dPBS, cells were permeabilized and blocked in 0.22 µm filter-sterilized 1X blocking buffer (10% normal goat serum (Thermo Scientific; 50197Z), 0.3% Triton X-100, 0.002% NaN_3_) for 1 hr at room temperature. Cells were incubated with the following primary antibodies diluted in blocking buffer overnight at 4°C: mouse anti-FLAG M2 (1:500; Sigma; F1804), rabbit anti-PEX14 (1:500; EMD Millipore; ABC142) and chicken anti-HSP60 (1:500; EnCor Biotechnology; CPCA-HSP60). Samples were washed 3X in PBS-T (0.3% Triton X-100) for 10 mins each, following by incubation with the following secondary antibodies: Goat anti-Mouse IgG (H+L) Cross-Adsorbed Secondary Antibody, Alexa Fluor 488 (1:500; Thermo Scientific; A-11001), Goat anti-Rabbit IgG (H+L) Cross-Adsorbed Secondary Antibody, Alexa Fluor 568 (1:500; Thermo Scientific; A-11011), Goat anti-Chicken IgY (H+L) Cross-Adsorbed Secondary Antibody, Alexa Fluor Plus 647 (1:500; Thermo Scientific; A32933) and counterstained with 1 µg/ml DAPI for 1 hr at room temperature. Samples were washed with 3X PBS-T prior to mounting on Superfrost Plus slides (Fisher Scientific; 12-550-15) with Fluoromount-G (SouthernBiotech; 0100-01). Samples were imaged on a Zeiss LSM 880 II Airyscan FAST confocal microscope using a 63X objective and an Airyscan module. ImageJ software (v2.9.0/1.53t) was used to analyze fluorescence intensity profiles of czi files.

### Liver RNA isolation and qPCR analysis

Frozen liver tissue (∼50-70 mg) was pulverized with a sterile metal bead in 1 mL TRIzol reagent (Thermo Fisher) by shaking with the TissueLyser II (Qiagen) (30/s frequency, 40 sec, 3 cycles). In between each cycle, samples cooled at 4°C for 5 mins. Samples were transferred to a new tube and mixed with 200 µL of cold chloroform. After inverting ten times, sample was centrifuged (12000 x *g*, 15 mins, 4°C). The upper phase was transferred to a fresh tube containing 500 µL of cold, molecular-grade IPA (Fisher Scientific) and precipitated at −80°C overnight. After centrifugation (12000 x *g*, 15 mins, 4°C), the supernatant was aspirated. RNA pellet was washed twice with 1 mL of cold 75% EtOH and centrifuged each time (7500 x *g*, 2 mins, 4°C). The supernatant was carefully aspirated and RNA pellets dried in a fume hood. Isolated RNA was dissolved in UltraPure water and quantified by NanoDrop. cDNA was synthesized from 2 µg RNA using the High-Capacity cDNA Reverse Transcriptase Kit (Thermo Scientific). 20 µL cDNA reactions were diluted 1:10 in UltraPure water. qPCR experiments were prepared in 384-well plates. Diluted cDNA was mixed with target primers and Applied Biosystems PowerUp SYBR Green Master Mix (Thermo Scientific) according to manufacturer instructions. Standard curves were generated by pooling cDNA samples. qPCR experiments were performed on the QuantStudio 5 instrument using default settings compatible with SYBR reagent. Gene expression was normalized to *Rps3* abundance.

### Statistics

All *in vitro* enzyme assay and cell culture data are expressed as mean -/+ SD. Animal experimental data are expressed as mean -/+ SEM. Statistical significance was calculated using the two-sided Student’s *t* test. Across the study, statistical significance depicted in data figures as asterisks are as follows: * = *P* < 0.05; ** = *P* < 0.005; *** = *P* < 0.0005. Two-way ANOVA metabolomic data was performed using the GraphPad Prism software (version 9.4.1).

## Acknowledgements

We would like to thank the Pagliarini Lab for their helpful feedback and discussion throughout the duration of this study. We are also grateful to the members of the Simcox Lab for their assistance with animal experiments. This work was supported by NIH awards R35 GM131795 and R01 R01DK098672 (D.J.P.), T32 GM008349 (E.H.R. and A.K.B.), SciMed Advanced Opportunity Fellowship (E.H.R.), and funds from the BJC Investigator Program (D.J.P.). This study leveraged the services of the Genome Engineering and Stem Cell Center at Washington University in St. Louis to generate CRISPR/Cas9 KO cells and of the Animal Models Core at the University of Wisconsin-Madison to establish whole body CRISPR/Cas9 KO mice. We thank the Washington University Center for Cellular Imaging for assistance with the preparation of cryo-EM experiments and data collection. We would like to thank Mingzhou Zhou and WuXi AppTec for their synthesis of custom compounds used in this study, and Bruce Palfey for generously sharing stocks of 5-deazaFAD. We also thank Vernon Anderson for his helpful discussions and insights on 4-HA metabolism and biochemistry.

## Author Contributions

E.H.R., A.K.B., and D.J.P. led the conception, design, and execution of this study and wrote this manuscript. E.H.R. and A.K.B. performed *in silico* analyses of sequence homology, cloned and purified recombinant enzymes, and performed *in vitro* enzyme assays. A.K.B., J.Z., and P.Y. performed cryo-EM experiments and structural analyses, and A.K.B. conducted structure-guided experiments. A.J.S. performed cell transfection and immunofluorescence microscopy *in vitro* experiments. E.H.R. performed lipid metabolism studies with cultured cells and extracted metabolites and lipids for LC-MS analysis. D.B.K., Z.N.B., and D.A.N. performed LC-MS metabolomics of acyl-CoAs. R.J. and J.A.S. developed lipidomic methods and measured 4-HAs *in vivo*. A.C. assisted with literature research and cloning. E.H.R., A.R.C, and J.A.S. maintained mouse colonies and performed mouse experiments. T.C. and B.F.P. provided consultation and assisted with cloning and purifying LvaE.

## Supplementary Information

Supplementary Table 1 – Plasmid information and primer sequences

Supplementary Table 2 – Fragmentation and retention time information for the identification and quantification of acyl-CoAs and 4-HAs by LC-MS

Supplementary Table 3 – Reagent and internal standard information

Supplementary Figure 1 – HADDOCK report of octanoyl-CoA docking to ACAD11 A* structure

---

**Extended Data Fig. 1.**
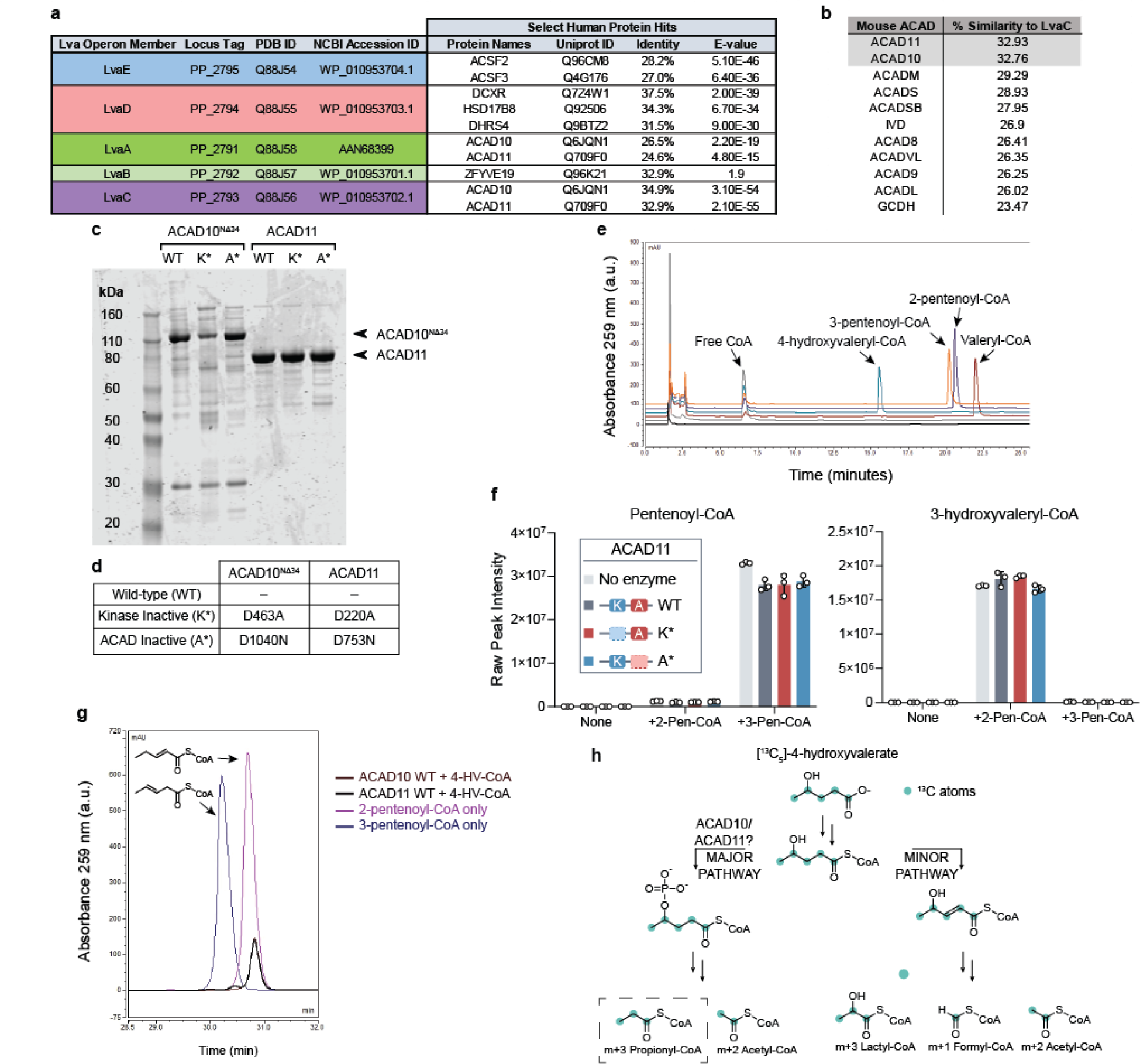
**(a)** Sequence identity of *lva* operon enzymes to human proteins as reported by UniProtKB database. **(b)** Sequence similarity of LvaC relative to mouse ACAD family according to pairwise alignments. **(c)** SDS-PAGE gel of recombinant mouse ACAD10^NΔ34^ and ACAD11 protein preparations from *P. pastoris* or RIPL *E. coli* orthogonal expression systems, respectively. WT = wild type; K* = Kinase inactive mutant; A* = ACAD inactive mutant. **(d)** Table describing the site-specific mutations to inactivate recombinant ACAD10^NΔ34^ and ACAD11 APH and ACAD catalytic domains. **(e)** HPLC chromatogram of acyl-CoA standards. **(f)** Assay testing the hydration activity of recombinant ACAD11. 2-pentenoyl-CoA (2-Pen-CoA) and 3-pentenoyl-CoA (3-Pen-CoA) were tested as substrates. Raw signal intensities of peaks corresponding to substrates and 3-hydroxyvaleryl-CoA product are depicted. Data represents mean -/+ SD (n = 3 technical replicates). **(g)** HPLC trace of pentenoyl-CoA isomers formed from 500 µM 4-HV-CoA by recombinant wild-type ACAD10^NΔ34^ and ACAD11 *in vitro* (brown and black traces, respectively) in comparison to LvaE-generated pentenoyl-CoA standards (dark purple and blue traces, respectively). **(h)** Expected labeling pattern of acyl-CoA catabolites derived from the catabolism of [^13^C_5_]-4-hydroxyvalerate (4-HV) by the “major” or “minor” pathways. M+3 propionyl-CoA labeling reflects catabolism through the 4-phosphovaleryl-CoA intermediate, presumably catalyzed by ACAD10/11.

**Extended Data Fig. 2.**
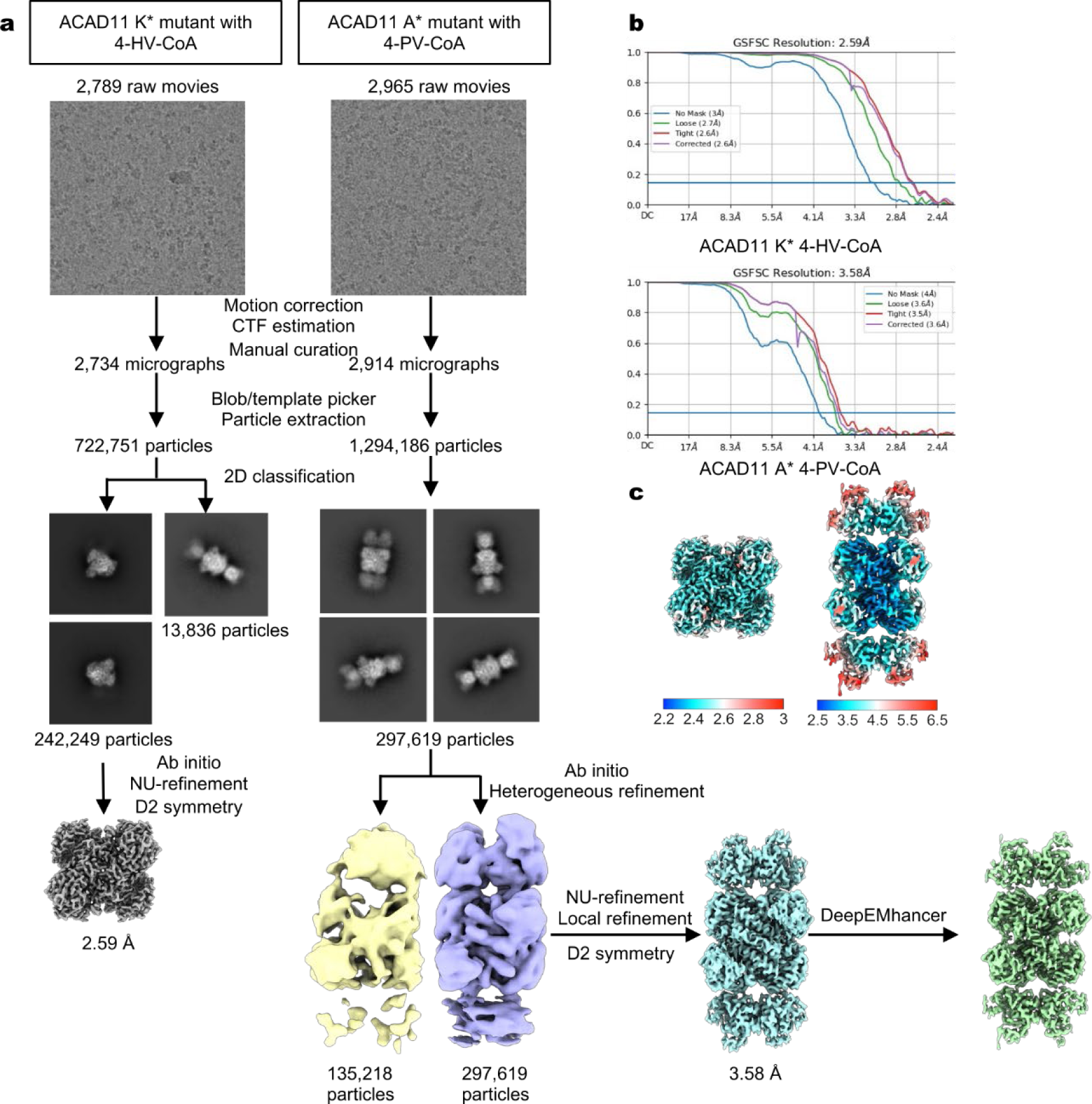
Cryo-EM analysis of the ACAD11 K* and A* mutants. **(a)** Data processing flowcharts. **(b)** FSC curves of non-uniform refinement of the ACAD11 K* mutant and local refinement of the ACAD11 A* mutant. **(c)** Local resolution of the final maps.

**Extended Data Fig. 3.**
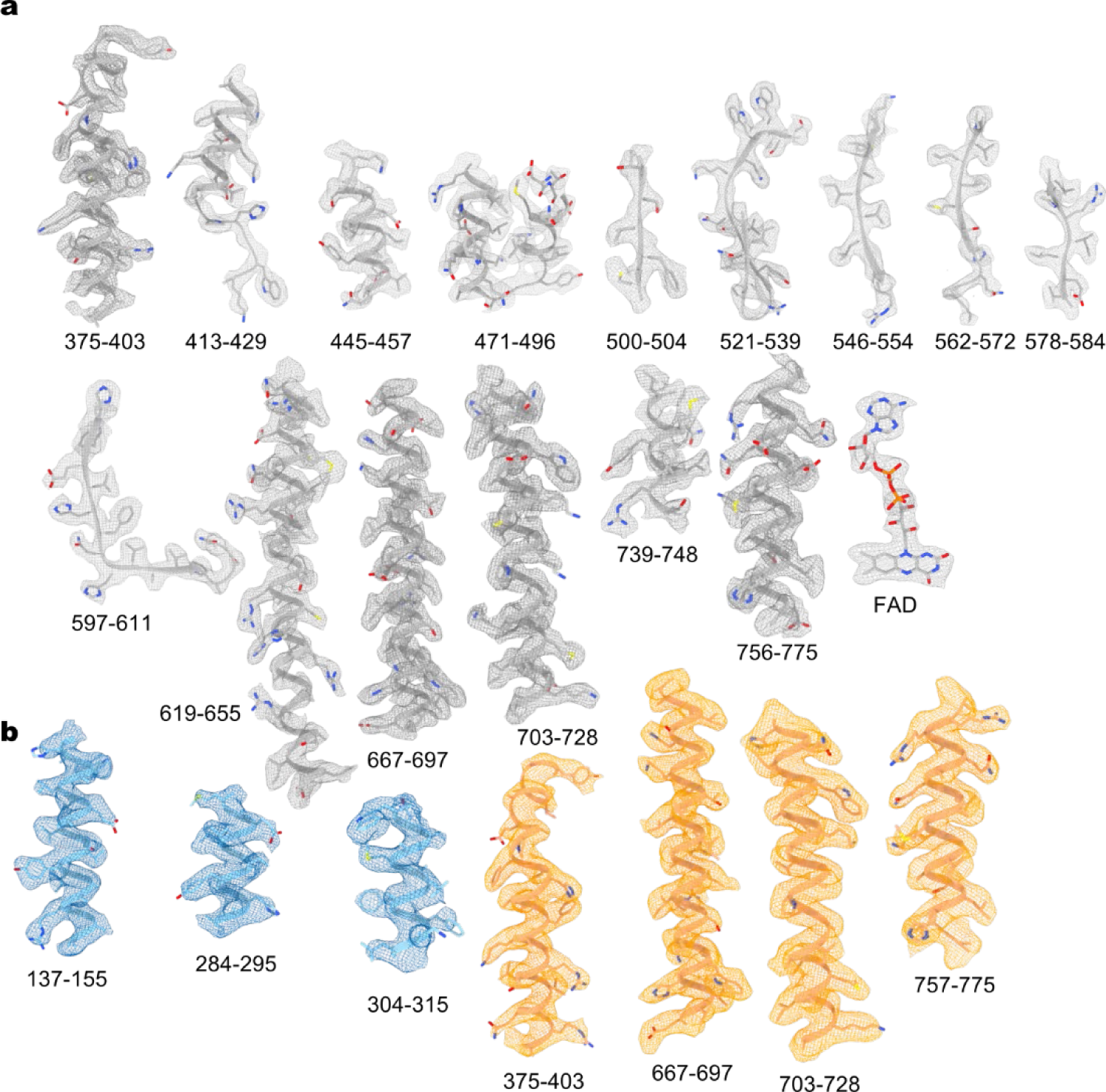
Representative cryo-EM densities of the ACAD11 K* mutant (**a**) and A* mutant (**b**).

**Extended Data Fig. 4.**
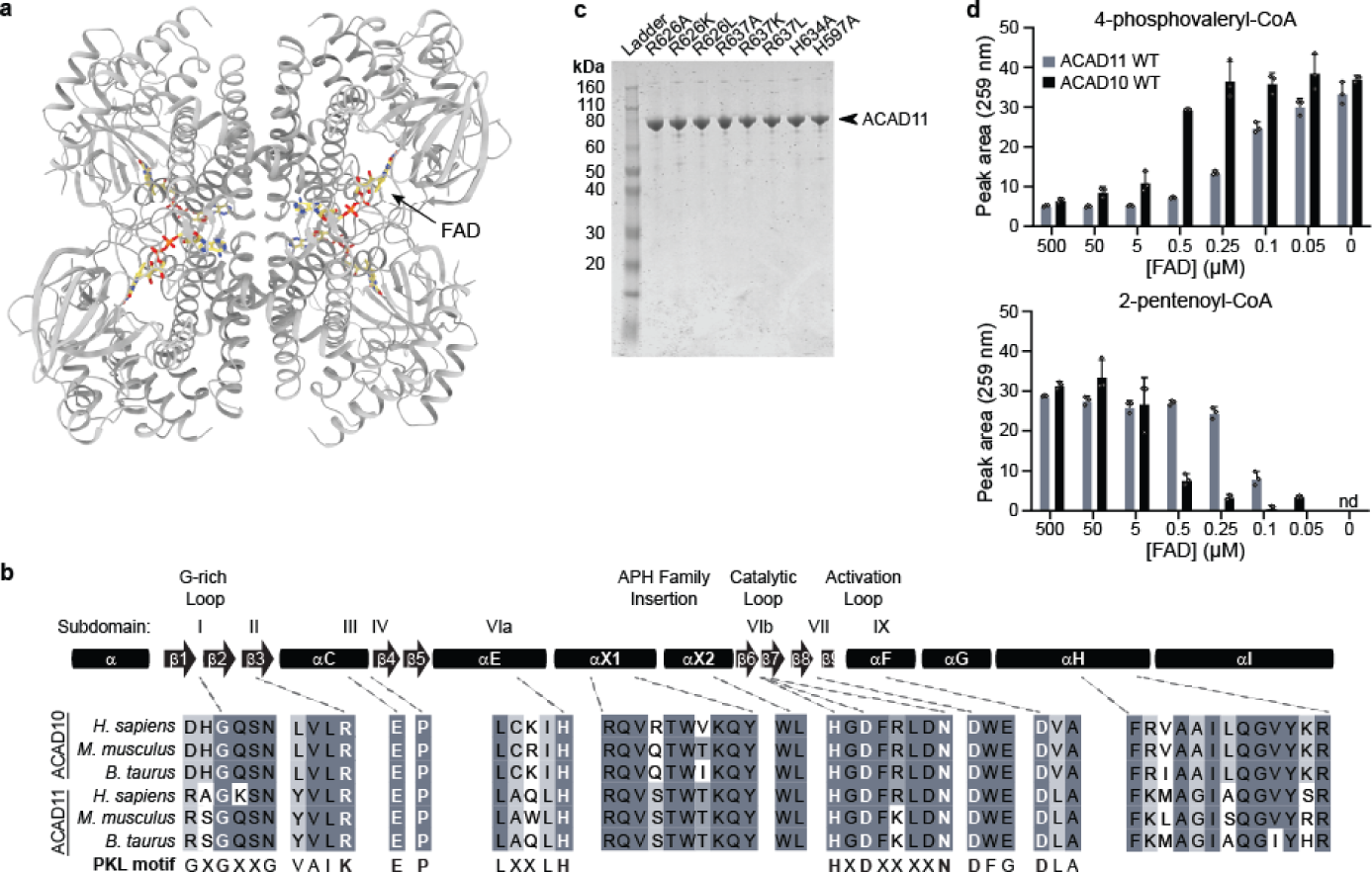
**(a)** Cryo-EM structure of the ACAD11 K* mutant with 4-HV-CoA. FAD molecules are highlighted in yellow. **(b)** Sequence alignment for key regions of the kinase domain. The overall canonical PKL subdomain structure is shown at the top. ACAD10/11 homologs from *H. sapiens*, *M. musculus*, and *B. taurus* are compared to PKL consensus motifs, with key conserved residues highlighted. **(c)** Coomassie-stained gels of recombiniant ACAD11 ACAD-domain mutants purified from *E. coli*. **(d)** UV-Vis HPLC quantification (259 nm) of 4-phosphovaleryl-CoA and 2-pentenoyl-CoA in reactions containing wild type ACAD10/11 and varying concentrations of FAD (mean -/+ SD; n = 3 technical replicates).

**Extended Data Fig. 5.**
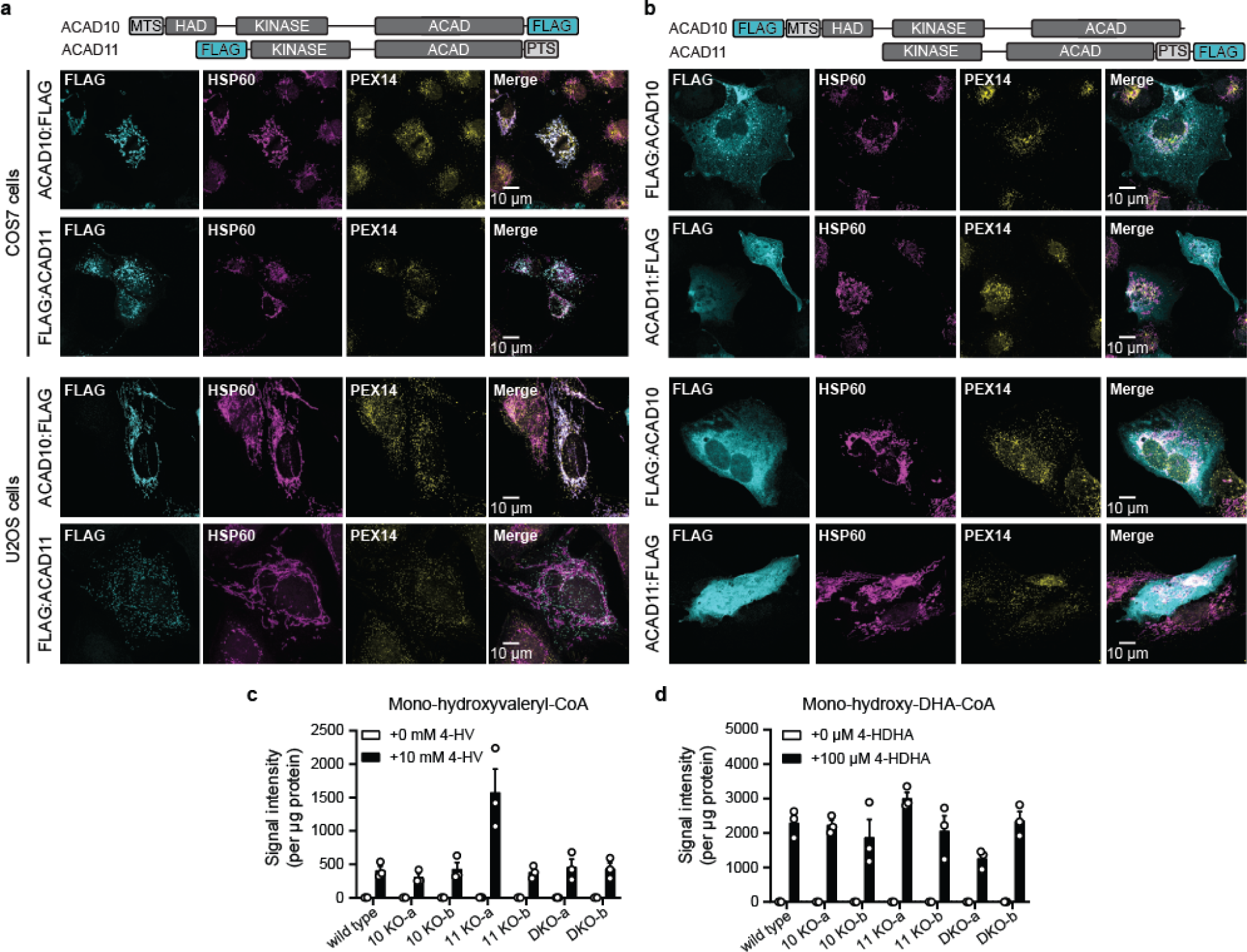
**(a)** Immunofluoresence imaging of overexpressed FLAG-tagged constructs in COS7 and U2OS cells at 10 µm scale. **(b)** Immunofluoresence imaging of overexpressed constructs with FLAG-tags swapped to opposite termini in COS7 and U2OS cells at 10 µm scale. **(c-d)** Peak intensity measurements likely corresponding to 4-hydroxyacyl-CoAs of exogenously delivered 4-HAs. Data represented as the mean signal intensity after protein normalization -/+ SD (n = 3 replicate experiments).

**Extended Data Fig. 6.**
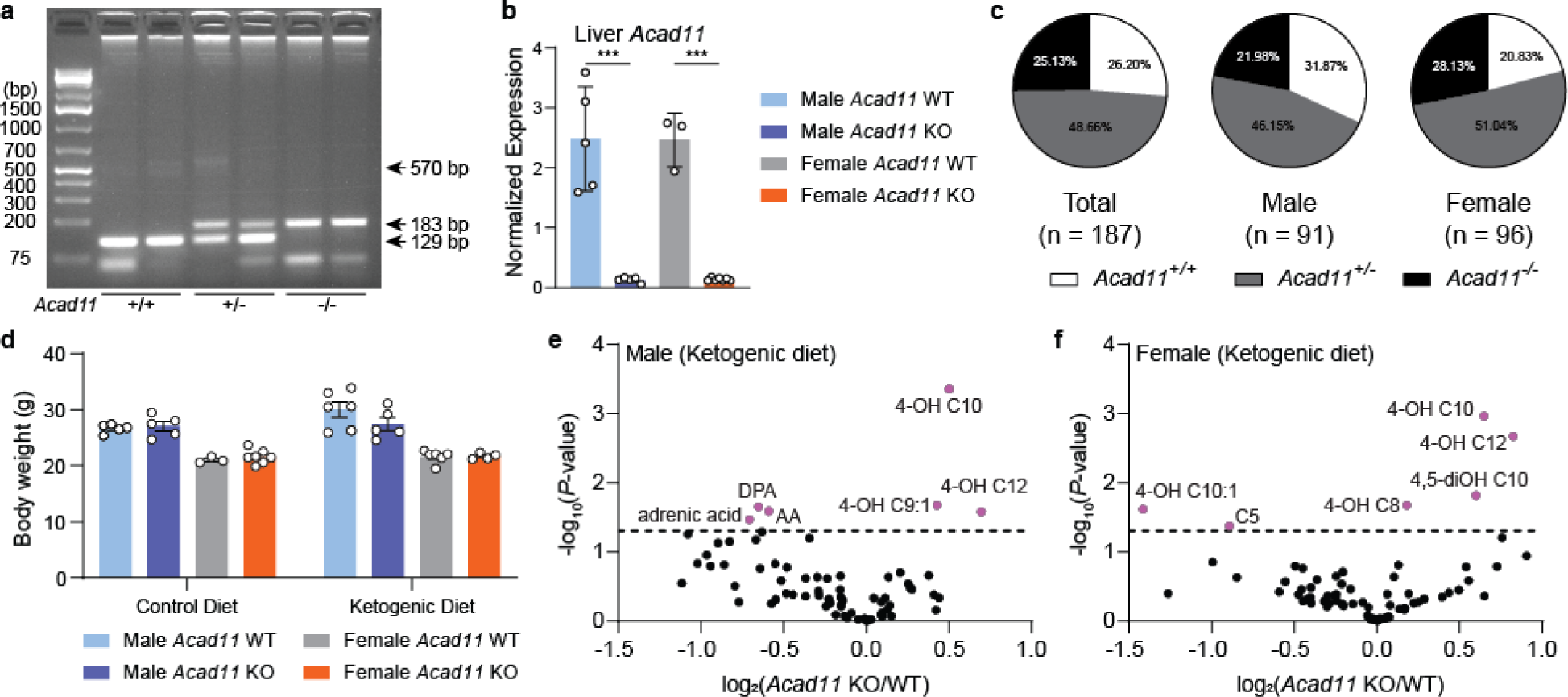
**(a)** Genotyping for *Acad11* KO mouse colony. **(b)** Normalized mRNA expression of *Acad11* in liver tissue. Data are normalized to *Rps3* levels. **(c)** Mendalian birth ratios of *Acad11* mice. **(d)** Body weight measurements (grams) of control diet- and ketogenic diet-fed mice. **(e-f)** Volcano plots of log_2_-transformed fold changes of 4-HAs and other targeted hydroxylated lipids relative to statistical significance after feeding on a ketogenic diet. Male and female KO mice were compared to respective littermate controls.

**Extended Data Table 1.** Cryo-EM data collection, refinement, and validation statistics.

|  | ACAD11 K* 4-HV-CoA<br>(EMD-42954)<br>(PDB 8V3U) | ACAD11 A* 4-PV-CoA<br>(EMD-42955)<br>(PDB 8V3V) |
| --- | --- | --- |
| **Data collection and processing** |  |  |
| Magnification | 59k | 59k |
| Voltage (kV) | 300 | 300 |
| Electron exposure (e<sup>-</sup>/Å<sup>2</sup>) | 54 | 54 |
| Defocus range (μm) | -1.0 to -2.4 | -1.0 to -2.4 |
| Pixel size (Å) | 1.1 | 1.1 |
| Symmetry imposed | D2 | D2 |
| Initial particle images (no.) | 722,751 | 1,294,186 |
| Final particle images (no.) | 242,249 | 297,619 |
| Map resolution (Å) | 2.6 | 3.6 |
| FSC threshold | 0.143 | 0.143 |
| Map resolution range (Å) | 2.2-3.0 | 2.5-6.5 |
| **Refinement** |  |  |
| Initial model used (PDB code) | AlphaFold and PDB: 2WBI | This study and AlphaFold |
| Model resolution (Å) | 2.7 | 3.7 |
| FSC threshold | 0.5 | 0.5 |
| Map sharpening *B* factor (Å<sup>2</sup>) | -137.8 | -144.7 |
| Model composition |  |  |
| Nonhydrogen atoms | 11,928 | 14,468 |
| Protein residues | 1,576 | 2,264 |
| Ligands | 4 | 4 |
| *B* factors (Å<sup>2</sup>) |  |  |
| Protein | 29.19 | 60.32 |
| Ligand | 16.77 | 78.76 |
| R.m.s. deviations |  |  |
| Bond lengths (Å) | 0.003 | 0.002 |
| Bond angles (°) | 0.461 | 0.447 |
| Validation |  |  |
| MolProbity score | 1.15 | 1.36 |
| Clash score | 3.55 | 4.66 |
| Poor rotamers (%) | 0.71 | 0.47 |
| Ramachandran plot |  |  |
| Favored (%) | 98.97 | 97.36 |
| Allowed (%) | 1.03 | 2.64 |
| Disallowed (%) | 0 | 0 |

